# MiR-9-5p protects from kidney fibrosis by metabolic reprogramming

**DOI:** 10.1101/667972

**Authors:** Marta Fierro-Fernández, Verónica Miguel, Laura Márquez-Expósito, Cristina Nuevo-Tapioles, J. Ignacio Herrero, Eva Blanco-Ruiz, Jessica Tituaña, Carolina Castillo, Pablo Cannata, María Monsalve, Marta Ruiz-Ortega, Ricardo Ramos, Santiago Lamas

## Abstract

MicroRNAs (miRNAs) regulate gene expression post-transcriptionally and control biological processes, including fibrogenesis. Kidney fibrosis remains a clinical challenge and miRNAs may represent a valid therapeutic avenue. We show that miR-9-5p protected from renal fibrosis in the mouse model of unilateral ureteral obstruction (UUO). This was reflected in reduced expression of pro-fibrotic markers, decreased number of infiltrating monocytes/macrophages and diminished tubular epithelial cell injury and transforming growth factor-beta 1 (TGF-β1)-dependent de-differentiation in human kidney proximal tubular (HKC-8) cells. RNA sequencing (RNA-Seq) studies in the UUO model revealed that this protection was mediated by a global shift in the expression profile of genes related to key metabolic pathways, including mitochondrial dysfunction, oxidative phosphorylation (OXPHOS), fatty acid oxidation (FAO) and glycolysis, preventing their UUO-dependent down-regulation. This effect was mirrored by a prevention in the TGF-β1-induced bioenergetics changes in HKC-8 cells. The expression of the FAO-related axis peroxisome proliferator-activated receptor gamma coactivator 1 alpha (PGC-1α)-peroxisome proliferator-activated receptor alpha (PPARα) was reduced by UUO, although preserved by the administration of miR-9-5p. We found that in mice null for the mitochondrial master regulator PGC-1α, miR-9-5p was unable to promote a protective effect in the UUO model. We propose that miR-9-5p elicits a protective response to chronic kidney injury and renal fibrosis by inducing reprogramming of the metabolic derangement and mitochondrial dysfunction affecting tubular epithelial cells.

## Introduction

The progression of chronic kidney disease (CKD) leads invariably to end-stage renal disease (ESRD), which is fatal without renal replacement therapy. Kidney fibrosis is the main hallmark of ESRD and where most progressive nephropathies converge. Histological manifestations of kidney fibrosis include the loss of capillary networks and the accumulation of fibrillary collagens, activated myofibroblasts and inflammatory cells (1, 2).

Recent studies have demonstrated a global defect in FAO leading to a compromise in the main source of energy for tubular epithelial cells (3). TGF-β1 is considered the master regulator of myofibroblast differentiation in fibrosis (4, 5) and acts as a key mediator of the reduction in FAO through Smad3-mediated repression of the *PPARGC1A* gene, which encodes for the peroxisome proliferator-activated receptor gamma coactivator 1 alpha (PGC-1α) protein (3). While there has been significant progress in the understanding of renal fibrogenesis (6-8), no effective therapy is available to prevent its progression or revert tissue damage. To date, many attempts have been made to stop or defer renal fibrosis, including the use of antihypertensive drugs (9), the blocking of TGF-β signaling pathways or the use of molecules with therapeutic effects in other organs (10). However, they have been scarcely effective (11-14) and thus there is urgent need to develop new therapeutic strategies.

miRNAs have emerged as critical regulators of gene expression through their action on post-transcriptional regulation. Their role in the progression and establishment of organ fibrosis has been reviewed elsewhere (15-17). In the kidney, miRNAs are not only indispensable for development and homeostasis, but also they are important players in pathophysiology(18-20). Several miRNAs have been shown to initiate or perpetuate cellular responses leading to fibrosis (21, 22). Notably, one of the best studied, miR-21, is closely related to fibrosis. Its expression is increased in both patient samples and animal models of CKD and acute kidney injury (AKI) (23-28). MiR-21 is up-regulated by TGF-β1 (29-31) and gene profiling identified several metabolic pathways, including alterations in lipid metabolism and FAO as pathogenic mediators of miR-21 effects (23, 32). Nevertheless, there are few reports identifying miRNAs with a protective action as ongoing therapeutic efforts are based on attempts to antagonize a specific miRNA, which given its multiplicity of targets may result in unwanted side effects (33).

In previous work we showed that treatment with miR-9-5p resulted in the prevention of fibrosis in several organs including the lung, peritoneum and skin (34, 35). We hypothesized that miR-9-5p could also be beneficial in renal fibrosis. In this work we present data showing that this miRNA protects the kidney from fibrosis by regulating metabolic pathways. We found that this effect is related to improved mitochondrial function and counteraction of TGF-β1 effects. Furthermore, we identify PGC-1α as an essential component in the protective action of miR-9-5p.

## Results

### Treatment with miR-9-5p prevents experimental renal fibrosis

The mouse model of UUO generates progressive renal fibrosis (36). To characterize the kinetics of miR-9-5p expression in the kidneys after UUO, we performed qRT-PCR analysis and we found that miR-9-5p expression was markedly enhanced at 5 days and remained elevated at 14 days after UUO (Figure 1A). To evaluate the role of miR-9-5p in renal fibrosis, lentiviral vectors expressing a scramble negative control construct (Lenti-SC) or miR-9-5p (Lenti-miR-9) were administered in mice 5 days before UUO. Retro-orbital injection of Lenti-miR-9 increased the expression levels of miR-9-5p up to 2-fold compared to control mice (Figure 1B). Histological parameters were evaluated in tissue samples from mice after lentiviral injection and exposed to UUO for 5 and 10 days. Lenti-SC treated animals showed significant tubulo-interstitial pathological changes 5 (Figure 1C, Figure 2A, upper panels) and 10 (Figure 1D, Figure 2A, lower panels) days after UUO. These consisted in tubular atrophy (Figure 1, C-E), tubular dilatation (Figure 1, C, D and F) and interstitial fibrosis (Figure 2, A-C). The extent of tubular atrophy and dilatation was markedly reduced by miR-9-5p 10 days after UUO (Figure 1, C-F). To determine the effect of miR-9-5p on fibrosis, sirius red staining was performed to quantify the collagen content in the kidneys. Evaluation of renal lesions by light microscopy showed increased collagen deposition in the interstitium after 5 (Figure 2A, upper panels, Figure 2B) and 10 (Figure 2A, lower panels, Figure 2C) days UUO in the obstructed kidneys of mice injected Lenti-SC. A significant protective effect of miR-9-5p was observed by a reduction of 20-40% in collagen deposition. Lenti-SC-treated mice showed increased extracellular matrix (ECM) protein levels, as indicated by increased type I collagen accumulation, after 10 days UUO (Figure 2D). Importantly, miR-9-5p prevented collagen deposition (Figure 2D). This was accompanied by a decrease in α1 type-1 collagen (*Col1α1*) gene expression after 5 (Figure 2E, left panel) and 10 (Figure 2E, right panel) days UUO in the obstructed kidneys of Lenti-miR-9 administered mice. Lenti-miR-9 administration also reduced the mRNA expression levels of fibronectin (*Fn1*) induced by UUO (Figure 2F). In the obstructed kidneys of Lenti-SC injected mice, an increase in proliferating tubulo-epithelial and tubulo-interstitial cells, detected by PCNA staining, was observed, an effect abrogated by miR-9-5p (Supplemental Figure 1A and B). These results support that miR-9-5p has a marked protective effect in kidney fibrosis promoted by UUO.

**Figure 1.**
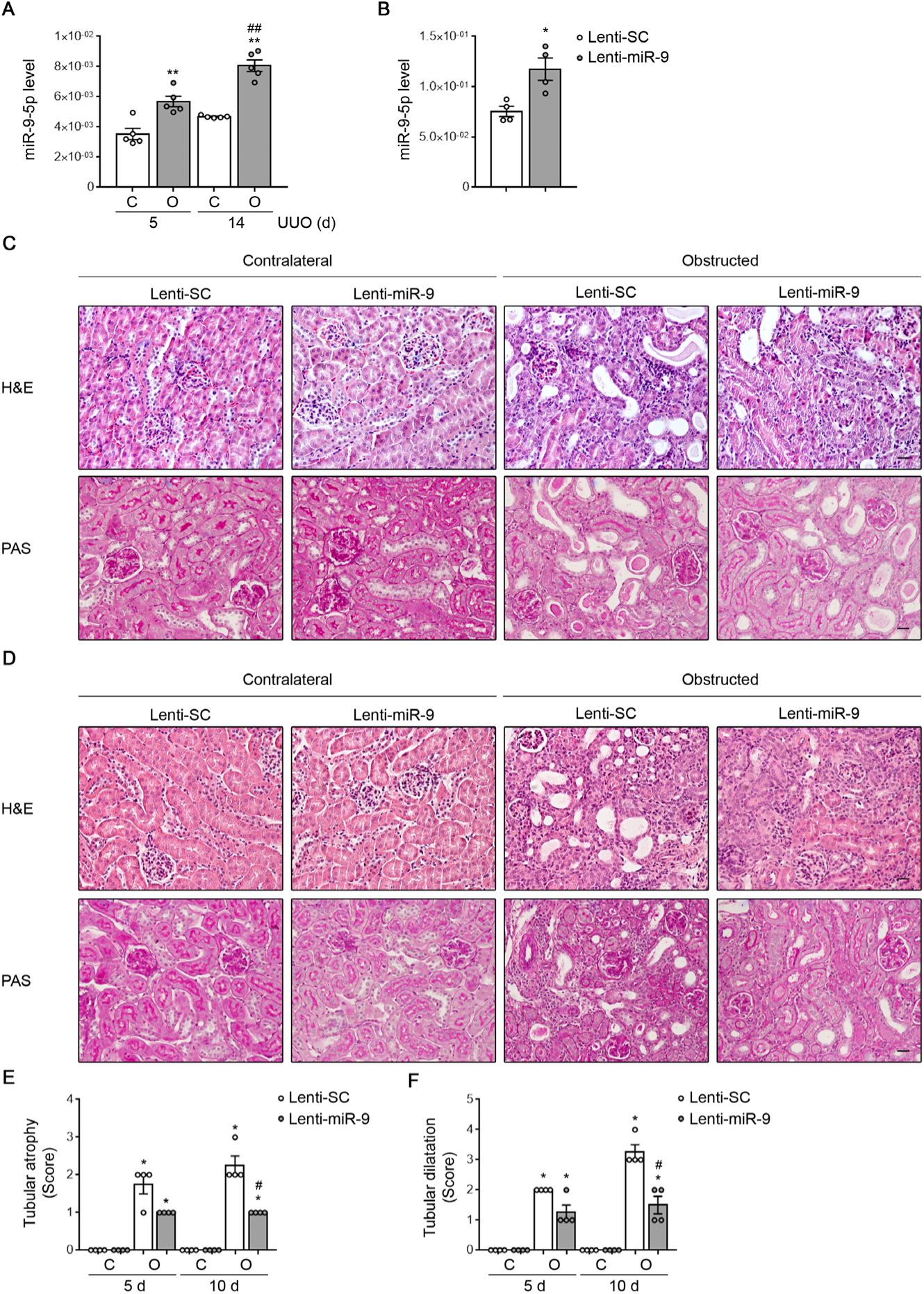
Over-expression of miR-9-5p protects from kidney fibrosis in the UUO mouse model. (**A**) qRT-PCR analysis of miR-9-5p expression levels in kidneys from mice after 5 and 14 days UUO. (**B**) Lentiviral particles (2×10^7^ ifu/mouse) containing miR-9-5p (Lenti-miR-9) or scramble miRNA (Lenti-SC) were administered to C57BL/6 mice for 5 days and miR-9-5p levels were determined by qRT-PCR in kidney tissue. *P < 0.05 compared to their corresponding Lenti-SC treated kidneys. (**C** and **D**) Representative microphotographs from one mouse per group of Hematoxylin and Eosin (H&E) (upper panels) and Periodic Acid Schiff (PAS) stainings (lower panels) from kidney sections of mice treated as described in (**B**) and subjected to UUO for 5 (**C**) and 10 (**D**) days. Microscopic images are representative of at least 4 mice per group. Scale bars: 50 µm. (**E** and **F**) Semi-quantitative determination (grade 0 to 4) of tubular atrophy (**E**) and tubular dilation (**F**) in kidney tissue samples from mice treated as described in (**C** and **D**). (**A**, **E** and **F**) C: contralateral, O: obstructed. *P < 0.05, **P < 0.01 compared to their corresponding contralateral kidneys. (**A**) ^##^P < 0.01, compared to 5 days obstructed kidneys. (**E** and **F**) ^#^P < 0.05, compared to obstructed kidneys in mice administered Lenti-SC. All statistical significance was determined using non-parametric two-tailed Mann-Whitney U test. Data are shown as the mean ± SEM (n = 4-5 mice per group).

**Figure 2.**
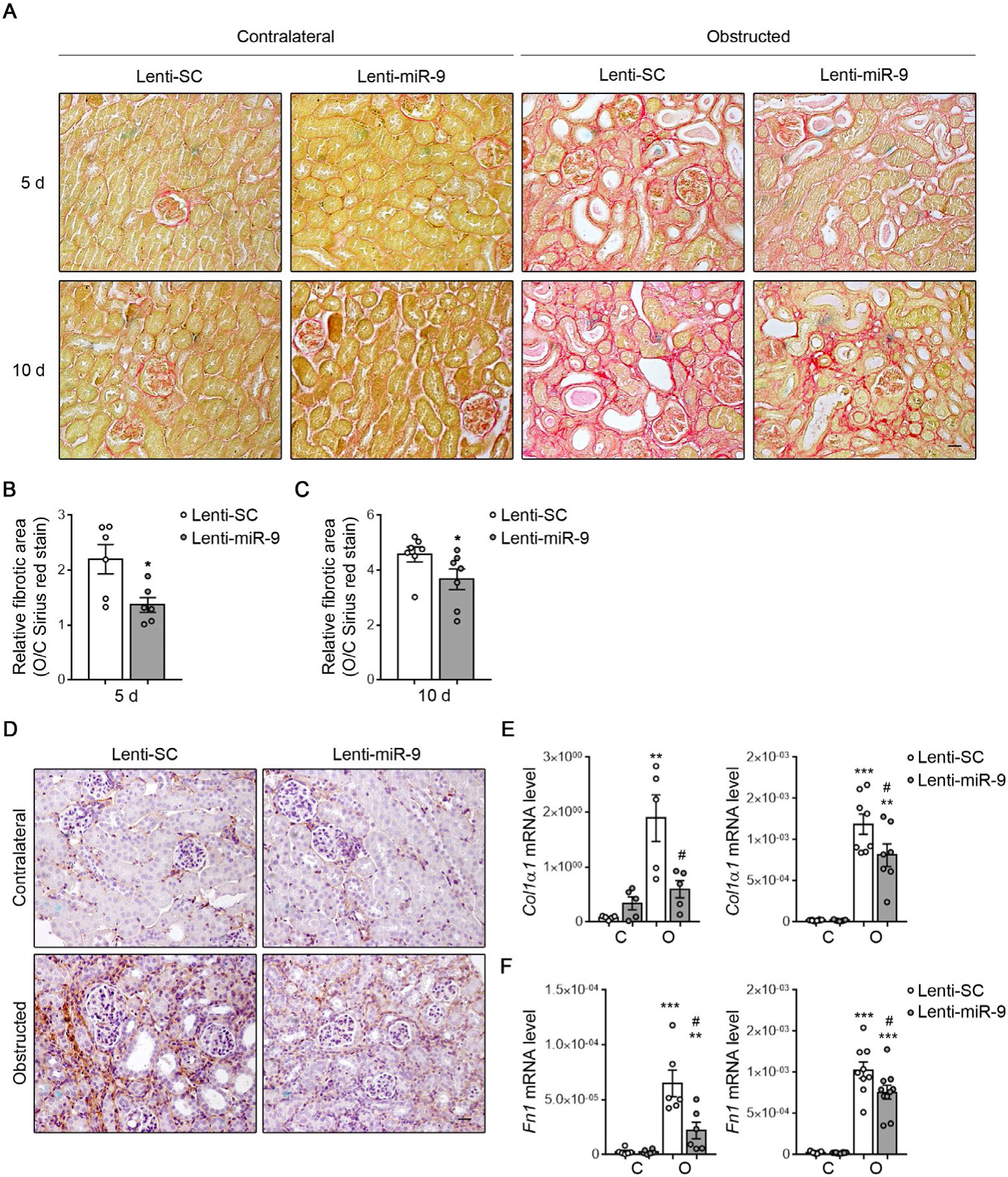
Increased miR-9-5p levels attenuate the deposition of extracellular matrix in the UUO mouse model. (**A**) C57BL/6 mice were injected with lentiviral particles (2×10^7^ ifu/mouse) containing miR-9-5p (Lenti-miR-9) or scramble miRNA (Lenti-SC) and 5 days later they were subjected to UUO. Representative microphotographs of one mouse per group of sirius red staining of kidneys from mice analyzed 5 (upper panels) or 10 (lower panels) days after UUO. (**B** and **C**) Quantification of sirius red staining, 5 (**B**) and 10 (**C**) days after UUO, was calculated as a ratio of the stained area over the total area. The fibrotic area is represented as the ratio of sirius red staining of obstructed (O) related to their corresponding contralateral (C) kidney (O/C) of mice treated as described in (**A**). *P < 0.05, compared to O/C kidneys in mice administered Lenti-SC. (**D**) Representative microphotographs of one mouse per group of the immunohistochemical staining for Type I Collagen from kidney sections of mice treated as described in (**A**) and subjected to UUO for 10 days. (**E** and **F**) mRNA levels of alpha 1 type-1 collagen (*Col1α1*) (**E**) and fibronectin (*Fn1*) (**F**) genes were determined by qRT-PCR in kidneys of mice treated as described in (**A**) and analyzed 5 (left panels) and 10 (right panels) days after UUO. **P < 0.01, ***P < 0.001 compared to their corresponding contralateral kidneys; ^#^P < 0.05, compared to obstructed kidneys in mice administered Lenti-SC. (**A** and **D**) Microscopic images are representative of at least 4 mice per group. Scale bars: 50 µm. (**B**, **C**, **E** and **F**) Statistical significance was determined using non-parametric two-tailed Mann-Whitney U test. Bar graphs data show the mean ± SEM (n = 5-10 mice per group).

### MiR-9-5p over-expression reduces myofibroblast formation after UUO

Myofibroblasts are a subset of activated fibroblasts that are considered the main cell type responsible for excessive synthesis and deposition of matrix proteins, especially Col1α1 and FN1, in kidney fibrosis (37-40). TGF-β1, one of the master regulators of fibrogenesis (41-43), induces the transformation of fibroblasts into myofibroblasts (5). Closely associated with the anti-fibrotic role of miR-9-5p, the extension of UUO-induced interstitial myofibroblasts, detected by α-SMA staining, was reduced by approximately 30% 10 days after UUO in Lenti-miR-9-injected mice (Figure 3, A and B). This observation was confirmed by analysis of total renal protein levels in kidneys from mice after lentivirus injection and subjected to UUO for 5 (Figure 3, C and D) and 10 days (Figure 3, E and F). A concomitant decrease in the *Acta2* gene (Figure 3G), which encodes for α-SMA, as well as in platelet-derived growth factor receptor beta (*Pdgfrb*) gene levels (Figure 3H), also considered part of the signature of the myofibroblast phenotypic transformation, was observed. Cytokine-mediated persistent epithelial injury leads to a maladaptive repair response with sustained tubular damage, chronic inflammation, proliferation of myofibroblasts and interstitial fibrosis (44, 45). To evaluate the effect of miR-9-5p on epithelial injury, we undertook flow cytometry analysis of epithelial cells from kidneys after 5 days UUO. We found that although the total number of epithelial cells (EPCAM^+^, CD45^−^) was clearly diminished in obstructed kidneys both in the absence and presence of miR-9-5p (Supplemental Figure 2, A and B), the proportion of damaged epithelial cells (CD24^+^) was slightly but significantly reduced in the obstructed kidneys from mice treated with Lenti-miR-9 (Supplemental Figure 2, A and C). To determine if miR-9-5p was involved in the pro-fibrogenic transformation of tubular epithelial cells by TGF-β1, HKC-8 cells were transfected with miR-9-5p and treated with TGF-β1 for different times. Increasing miR-9-5p levels significantly prevented the TGF-β1-reduced expression of *CDH1*, which encodes the epithelial cadherin (E-cadherin), and decreased the TGF-β1-induced expression of *ACTA2*, *COL1α1* and *FN1* (Supplemental Figure 3A). Similarly, over-expression of miR-9-5p strongly reduced α-SMA (Supplemental Figure 3, B and C) and FN1 protein abundance (Supplemental Figure 3, D and E). Consistently, over-expression of miR-9-5p attenuated SMAD2 phosphorylation after TGF-β1 stimulation (Supplemental Figure 3, F and G), suggesting that the inhibitory effect of miR-9-5p on the pro-fibrogenic transformation of HKC-8 cells was, at least in part, mediated by regulating TGF-β1-mediated signals.

**Figure 3.**
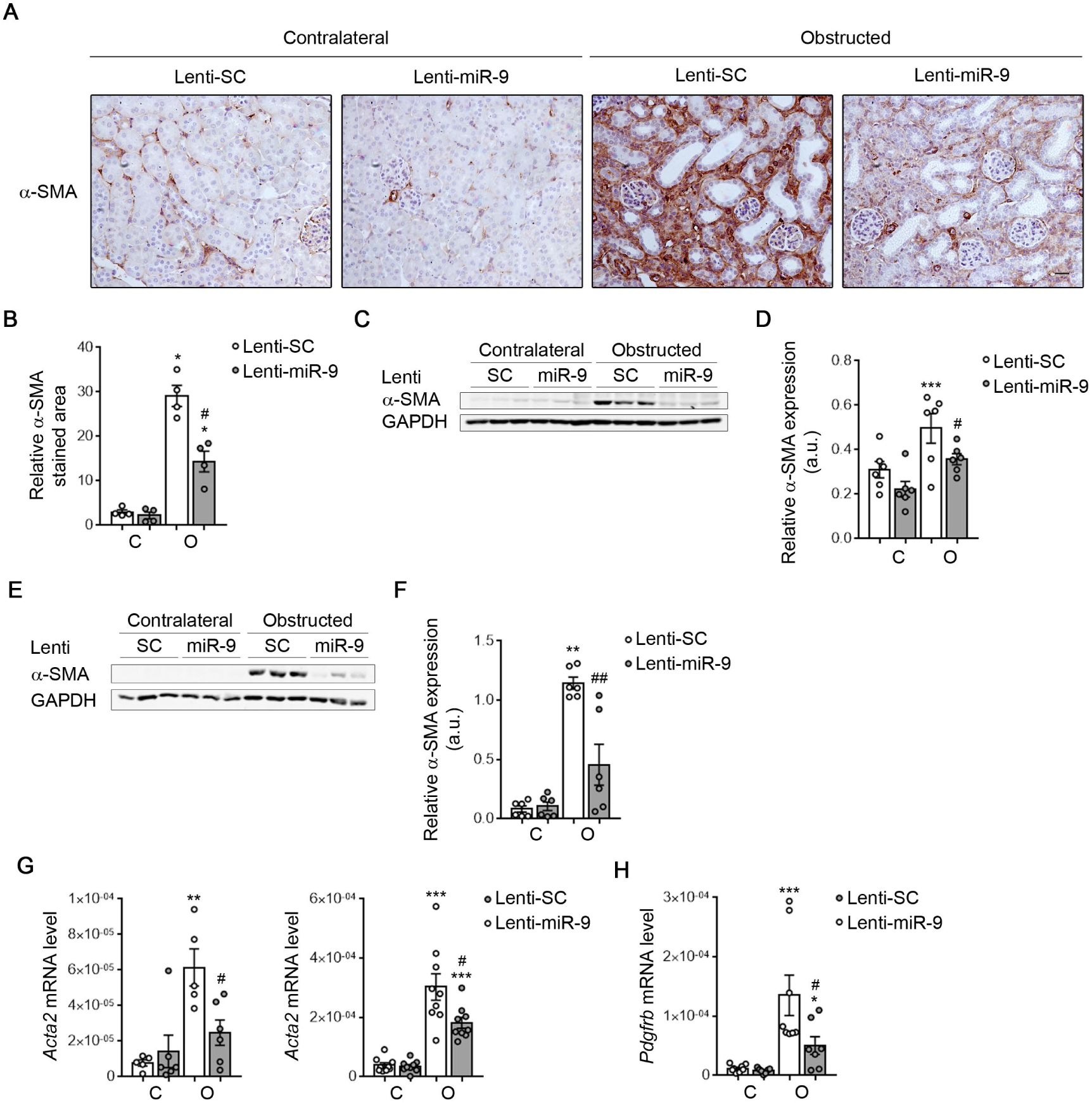
Up-regulation of miR-9-5p reduces the expression of fibrotic and myofibroblast markers after UUO. (**A**) Representative microphotographs of one mouse per group of immunohistochemical staining for alpha-smooth muscle actin (α-SMA) from kidney sections of C57BL/6 mice injected with lentiviral particles (2×10^7^ ifu/mouse) carrying miR-9-5p (Lenti-miR-9) or scramble miRNA (Lenti-SC) for 5 days. Kidneys samples were analyzed 10 days after UUO. Microscopic images are representative of at least 4 mice per group. Scale bar: 50 µm. (**B**) Bar graph represents the quantification of α-SMA positive stained area relative to the total analyzed area in the contralateral (C) and the obstructed kidneys (O) from mice treated as described in (**A**). (**C** and **E**) Mice were treated as shown in (**A**) and UUO was performed for 5 (**C**) and 10 (**E**) days. Representative western blots of α-SMA protein levels in kidneys from 3 individual mice per group are shown. (**D** and **F**) Bar graphs represent densitometric values (arbitrary units, a.u.) of the α-SMA expression. (**G**) mRNA levels of *Acta2* gene were determined by qRT-PCR in kidneys from mice treated as described in (**A**) and analyzed 5 (left panel) and 10 (right panel) days after UUO. (**H**) mRNA level of platelet-derived growth factor receptor beta (*Pdgfrb*) gene was determined by qRT-PCR in mice treated as described in (**A**) and analyzed 10 days after UUO. (**C**-**F**) Glyceraldehyde-3-phosphate dehydrogenase (GAPDH) was used for normalization purposes. (**B**, **D**, and **F**-**H**) Bar graphs represent the mean ± SEM (n = 4-9 mice per group). *P < 0.05, **P < 0.01, ***P < 0.001 compared to their corresponding contralateral kidneys; ^#^P < 0.05, ^##^P < 0.01 compared to obstructed kidneys in mice administered Lenti-SC, non-parametric two-tailed Mann-Whitney U test.

### Increased levels of miR-9-5p reduce the monocyte/macrophage population in UUO kidneys

The accumulation of macrophages correlates with renal fibrosis in human kidney disease (46). As shown in Figure 4, A and B, an increased infiltration of monocytes/macrophages, (F4/80^+^ cells) was observed in obstructed kidneys from Lenti-SC-treated mice 5 days after UUO. Over-expression of miR-9-5p significantly reduced the area of infiltrating cells in obstructed kidneys. Flow cytometry analysis revealed that UUO significantly increased the number of monocytes/macrophages (F4/80^+^, CD45^+^) in obstructed kidneys, an effect that was prevented by miR-9-5p (Figure 4, C and D), confirming the histological observations. Overall, these data suggest that miR-9-5p may protect from fibrosis by targeting pro-inflammatory cells responsible for tubular apoptosis and fibroblast transformation.

**Figure 4.**
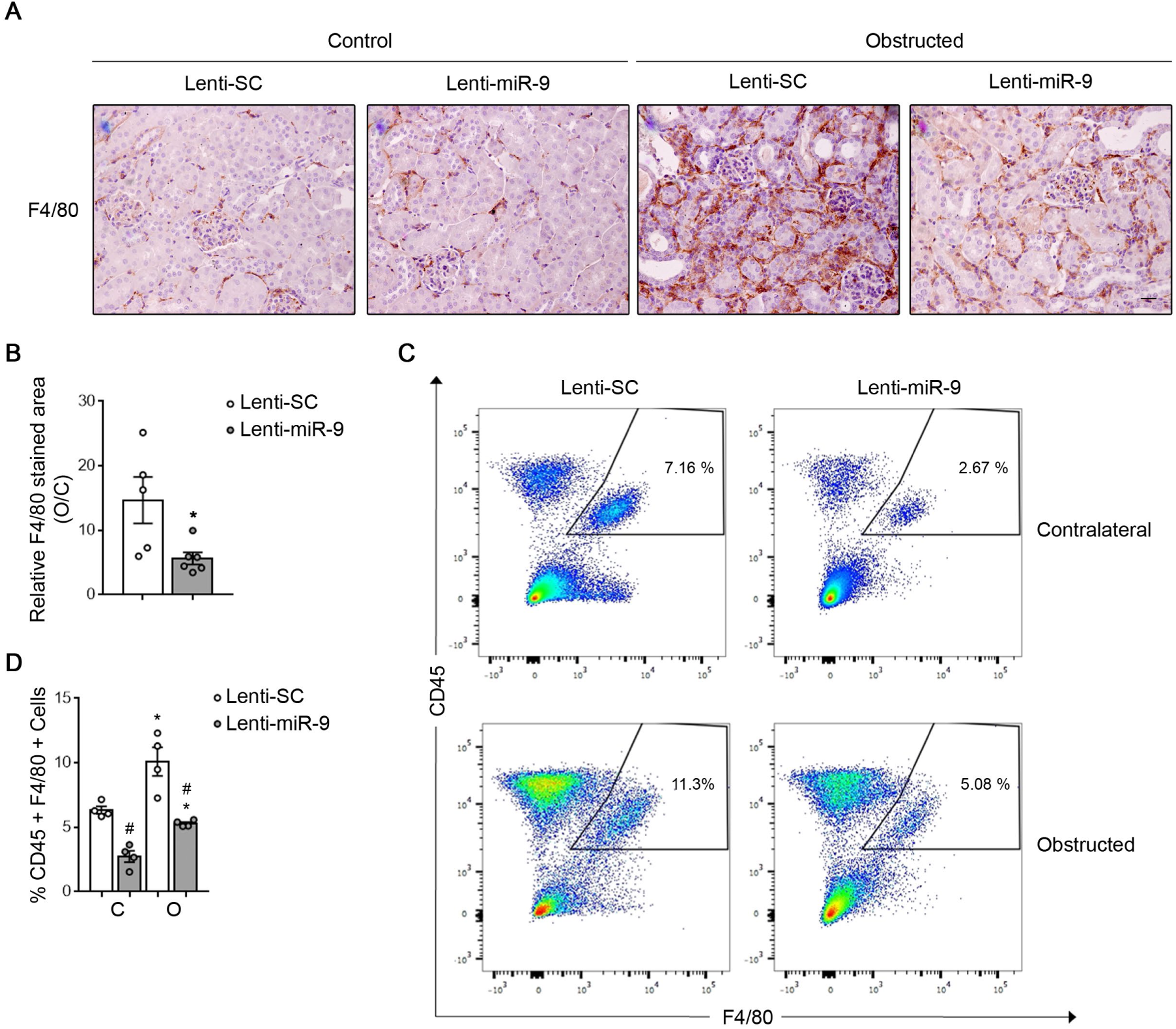
Administration of miR-9-5p reduces macrophage infiltration in UUO kidneys. (**A**) C57BL/6 mice were injected lentiviral particles (2×10^7^ ifu/mouse) containing miR-9-5p (Lenti-miR-9) or scramble miRNA (Lenti-SC) 5 days before UUO and analyzed 5 days after the procedure. Representative micrographs of one mouse per group showing the expression of F4/80 in kidney sections of mice treated as described above. Scale bar = 50 µm. (**B**) Bar graph represents the quantification of the ratio of F4/80 positive stained area relative to the total analyzed area in the obstructed (O) in relation to the corresponding contralateral (C) kidney (O/C) in mice treated as described in (**A**) (n = 5-6 mice per group). *P < 0.05, compared to O/C kidneys from mice administered Lenti-SC, non-parametric two-tailed Mann-Whitney U test. (**C**) Examples of multiparameter flow cytometry dot plots showing the expression of CD45, a panleukocyte antigen also present in hematopoietic cells, monocytes and macrophages, and F4/80, a surface marker expressed by monocytes and macrophages, in kidney cells from one mouse per group treated as described in (**A**). Numbers in quadrants indicate cell proportions in percent of cells that co-express both markers. Dot plots shown are representative of 4 mice per group. (**D**) Bar graphs represent the % of kidney cells co-expressing CD45 and F4/80 markers. Data represent mean ± SEM (n = 4 mice per group). *P < 0.05 compared to their corresponding contralateral kidneys; ^#^P < 0.05, compared to kidneys from mice administered Lenti-SC, non-parametric two-tailed Mann-Whitney U test.

### Transcriptomic analysis indicates that miR-9-5p regulates metabolic pathways in the fibrotic kidney

To understand the mechanisms underlying the protective action of miR-9-5p on renal fibrosis we performed RNA-Seq analysis to compare the global transcriptome of whole kidneys in mice injected Lenti-miR-9 or Lenti-SC for 5 days and subjected to UUO for 5 days. For the data analysis, kidney samples were classified in four groups (Figure 5A). After appropriate processing (Figure 5B), a total of 13,232 genes were considered for analysis. Only genes with a Q-value ˂ 0.01 were considered significantly different (Supplemental Figure 4A). The subsets of genes and their intersections among the experimental groups are depicted in Figure 5c and listed in Supplemental Table 1. Those genes that significantly changed in comparisons of 4 vs 2 and in 2 vs 1 groups were chosen for further analysis (highlighted in red numbers in Figure 5C). Almost all the genes, 99 %, were regulated in opposite directions by UUO and by miR-9-5p (Figure 5D, Supplemental Table 2). In-depth pattern analysis using two different bioinformatics resources, DAVID and iPathwayGuide softwares, identified differential regulation of pathways related to metabolism, OXPHOS, tricarboxylic acid (TCA) cycle and glycolysis (Supplemental Figure 4B, Supplemental Figure 5A, gray bars). Both *in silico* analyses indicated that metabolic pathways were the most prevalent among those significantly induced by miR-9-5p in the obstructed kidneys. Gene ontology analysis showed that oxidation-reduction, metabolic, ATP biosynthetic and lipid metabolism processes were among the biological processes (BPs) that changed more significantly (Supplemental Figure 5B, gray bars). In addition, BPs closely related to fibrosis were also modified by miR-9-5p (Supplemental Figure 5B, red bars). In fact, miR-9-5p prevented the increased expression of key pro-fibrotic genes such as *Acta2*, *Col1α1*, *Ctgf*, *Fn1*, *Pdgfrb*, and *Tgfb1* and its receptors among others (Supplemental Figure 6A), confirming the important role of this miRNA in renal fibrogenesis. TaqMan gene expression assays were confirmatory (Supplemental Figure 6B). Importantly, as we described in the context of lung fibrosis (34), miR-9-5p reduced the expression of transforming growth factor beta receptor II (*Tgfbr2*) and NADPH-oxidase 4 (*Nox4*) (Supplemental Figure 6, C and D), suggesting that both genes were also relevant targets for miR-9-5p in kidney fibrosis. Of note, RNA-Seq and Taqman expression analysis of inflammation-related genes showed that miR-9-5p blunted the increase of those induced by UUO (Supplemental Figure 6, E and F). Of interest, miR-9-5p suppressed the expression of the *Adgre1* gene, which encodes the F4/80 antigen, in consistence with its inhibitory effects on monocyte/macrophage infiltration (Figure 4). Taken together, these data suggest that miR-9-5p modifies a major portion of the genes related to metabolic pathways playing a role in kidney fibrosis (3, 32, 47).

**Figure 5.**
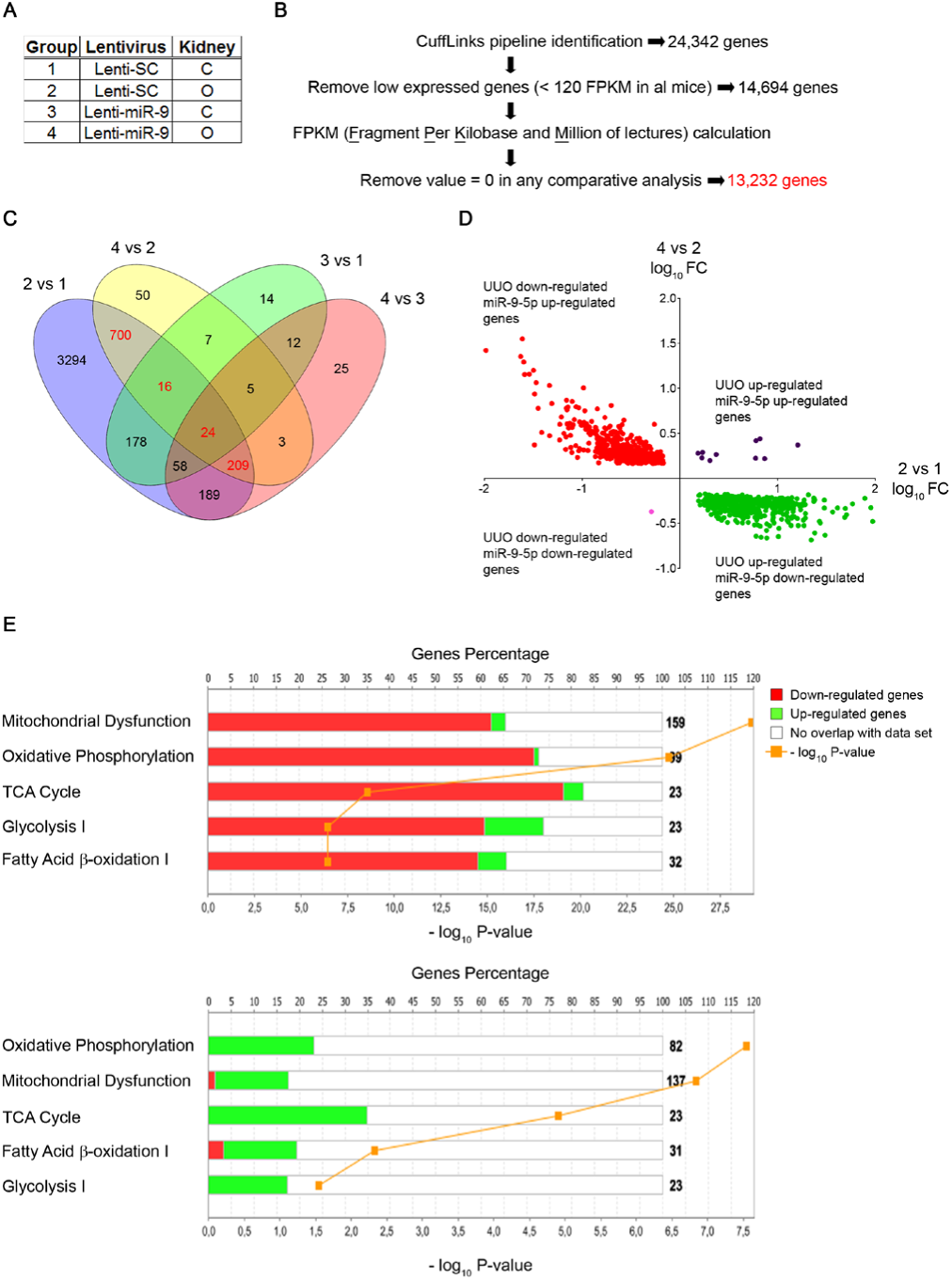
RNA-Seq analysis of kidneys from mice treated with Lenti-miR-5p and subjected to 5 days UUO. Lentiviral particles (2×10^7^ ifu/mouse) containing miR-9-5p (Lenti-miR-9) or scramble miRNA (Lenti-SC) were injected to C57BL/6 mice and 5 days later they were subjected to UUO for 5 days (n = 3 mice per group). (**A**) Summary of the analyzed kidneys groups. C: contralateral kidney; O: obstructed kidney. (**B**) Work flow summary of the data analysis and the number of genes resulting after corrections performed in each step. 13,232 genes were selected for further analysis. (**C**) Venn diagram showing the number of significant genes (Q-value ˂ 0.01) unique and overlapping between the datasets derived from all the possible comparisons among the groups shown in (**A**). The list of genes generated for this figure is shown in Supplemental Table 1. Numbers colored in red correspond to those genes whose expression changed after UUO and are selectively modified by miR-9-5p. (**D**) Dot plot graphic of log10 Fold Change (log10 FC) in gene expression for obstructed compared to control kidney from Lenti-SC treated mice (2 vs 1) (x-axis) represented against log10 FC in gene expression for obstructed kidney from Lenti-miR-9-treated mice compared to obstructed kidney from Lenti-SC treated mice (4 vs 2) (y-axis). Each dot represents an individual gene. The log10 FC from these genes and their corresponding P-value are listed in Supplemental Table 2. Explanatory legend of the dot colors is shown in the corresponding quadrant. (**E**) Differentially expressed genes associated with metabolic pathways in the Ingenuity Pathway Analysis (IPA) corresponding to the comparisons 2 vs 1 (upper panel, listed in Supplemental Table 3) and 4 vs 2 (lower panel, listed in Supplemental Table 4). The stacked bar charts display the percentage of genes (y-axis) distributed according to the direction of gene regulation changes in each canonical pathway (x-axis). Genes that were up-regulated in our dataset are shown in green, those that were down-regulated are shown in red and those found in the IPA reference gene set that had no overlap with our dataset are represented in white. The numerical value at the right of each canonical pathway bar indicates the total number of genes present in the IPA Knowledge data base for that pathway. The yellow line represents the significance of the directional change (–log10 P-value), calculated by adjusting the right-tailed Fisher’s Exact t-test P-values using the Benjamini-Hochberg method, of affected genes relative to the total number of genes in a pathway.

### MiR-9-5p reprograms changes in metabolic pathways induced by UUO

Ingenuity Pathway Analyses (IPA) indicated that, of the metabolism-related genes changed by miR-9-5p, the great majority were linked with mitochondrial dysfunction, OXPHOS, TCA cycle, glycolysis and FAO. These genes, commonly down-regulated by UUO (2 vs 1 comparison, Figure 5E, upper panel, red bars, Supplemental Table 3), were up-regulated in the obstructed kidney in response to miR-9-5p (4 vs 2 comparison, Figure 5E, lower panel, green bars, Supplemental Table 4). In addition, miR-9-5p also counteracted the down-regulation of the studied OXPHOS-related genes, including electron transport chain (ETC)- and ATP synthase-related genes, induced by UUO (Figure 6, A and B). Studies with Taqman probes were confirmatory (Figure 6C). A similar expression pattern was observed in glycolysis- and in TCA cycle-related genes (Figure 6D, Supplemental Figure 6G). Data obtained with the Taqman probes were also consistent (Figure 6E, Supplemental Figure 6H). Decreased levels of FAO are closely related to fibrotic damage in the kidney (3). We found that miR-9-5p was able to prevent the down-regulation of FAO-related genes 5 days after UUO (Figure 6F). This effect was corroborated by Taqman analysis (Figure 6G). In keeping, miR-9-5p also prevented the abrogation of the enzyme carnitine palmitoyl-transferase 1a (CPT1A), the rate-limiting enzyme in FAO (48), after 5 (Figure 6, H and I) and 10 days (Figure 6, J and K) UUO. Consistently, miR-9-5p restored mRNA levels of *CPT1a* reduced by UUO (Figure 6, L and M). These data support that miR-9-5p specifically reprograms metabolic pathways and counteracts the global derangement in metabolic dysfunction promoted by renal fibrosis.

**Figure 6.**
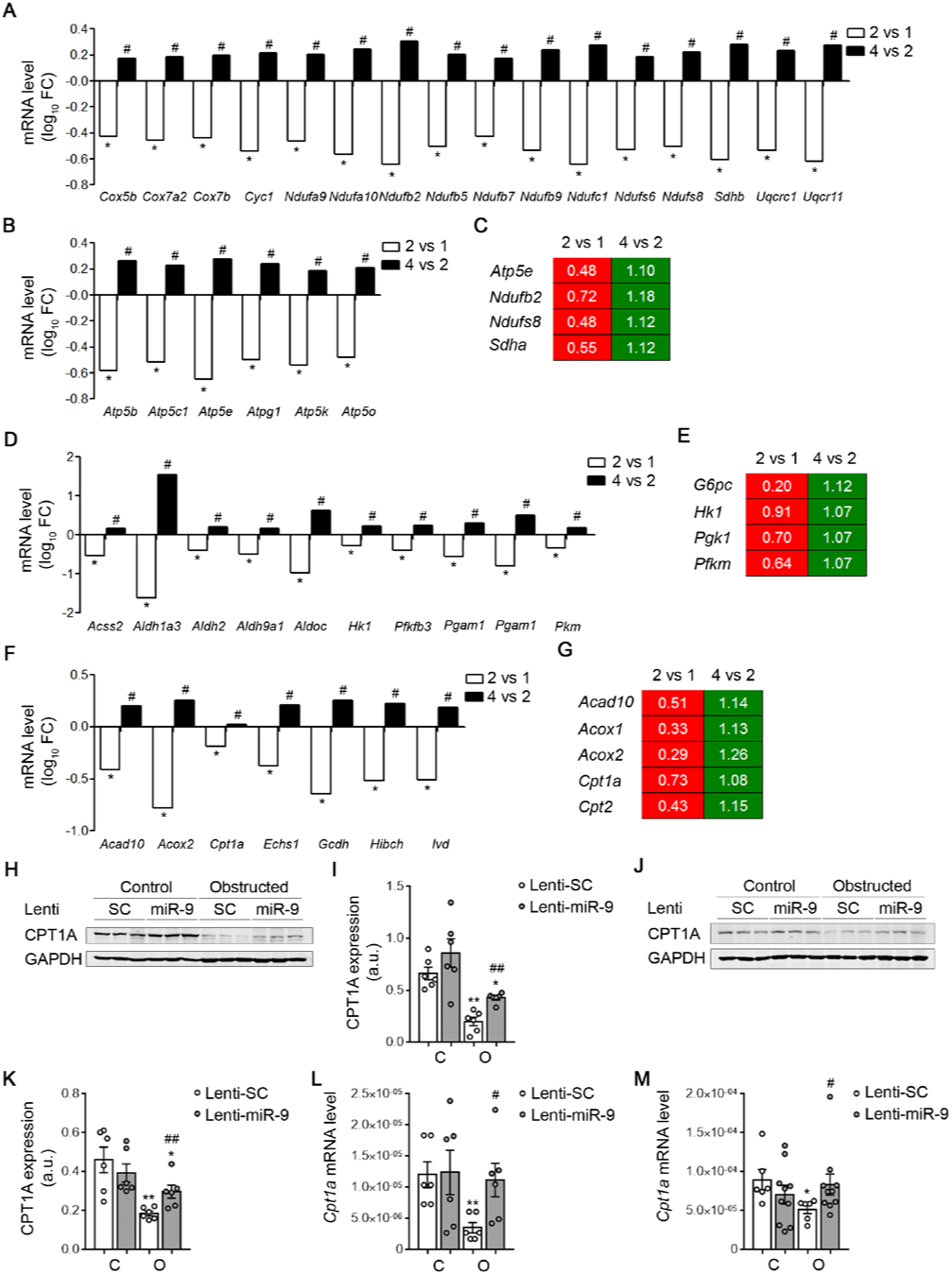
Administration of miR-9-5p reprograms the changes in metabolism-related genes induced by UUO. (**A**-**G**) Analysis of gene expression in kidney samples from C57BL/6 mice injected with lentiviral particles (2×10^7^ ifu/mouse) containing miR-9-5p (Lenti-miR-9) or scramble miRNA (Lenti-SC) 5 days before UUO and analyzed 5 days after the procedure. (**A** and **B**) RNA-Seq data pertaining to oxidative phosphorylation (OXPHOS)-related genes including electron transport chain (ETC)-related (**A**) and ATP synthase-related genes (**B**). (**C**) Gene expression heatmap generated by using qRT-PCR data of selected OXPHOS-related genes. (**D**) RNA-Seq data corresponding to glycolysis-related genes. (**E**) Gene expression heatmap generated by using qRT-PCR data of selected glucose utilization-related genes. (**F**) RNA-Seq data pertaining to β-oxidation of fatty acids (FAO)-related genes. (**G**) Gene expression heatmap generated by using qRT-PCR data of selected FAO-related genes. (**A**-**G**) Data correspond to comparisons 2 vs 1 and 4 vs 2 (groups described in Figure 5A). (**A**, **B**, **D** and **F**) RNA-seq data are represented as the log10 Fold Change (FC) (n = 3 mice per group), *Q < 0.01 compared to their corresponding contralateral kidneys; ^#^Q < 0.01 compared to obstructed kidneys in mice administered Lenti-SC, multiple hypothesis testing with the Benjamini– Hochberg FDR algorithm. (**C**, **E** and **G**) mRNA levels were analyzed by using Taqman qPCR probes (n = 6 mice per group). The numbers in each box indicate for each gene the mean of the fold change expression obtained from 6 mice per group. Red indicates that the gene was down-regulated and green that the gene was up-regulated. (**H** and **J**) Representative western blots of carnitine palmitoyltransferase 1A (CPT1A) protein levels in contralateral (C) and obstructed (O) kidneys from 3 individual mice per group injected as described above and subjected to UUO for 5 (**H**) and 10 days (**J**). (**I** and **K**) Bar graphs represent densitometric values (arbitrary units, a.u.) of CPT1A expression. (**H**-**K**) Glyceraldehide-3-phosphate dehydrogenase (GAPDH) was used for normalization purposes. (**L** and **M**) mRNA levels of *Cpt1a* gene in kidneys from mice injected as described in (**A**) and subjected to UUO for 5 (**L**) and 10 days (**M**) were analyzed by qRT-PCR. (**I** and **K**-**M**) Data are shown as the mean ± SEM (n = 6-10 mice per group), *P < 0.05, **P < 0.01 compared to their corresponding contralateral kidneys; ^#^P < 0.05, ^##^P < 0.01 compared to obstructed kidneys in mice administered Lenti-SC, non-parametric two-tailed Mann-Whitney U test.

### Over-expression of miR-9-5p recovers mitochondrial function in TGF-β1-treated HKC-8 cells

To gain insight into the metabolic consequences associated with the administration of miR-9-5p we undertook studies in HKC-8 cells and TGF-β1 was used as the model cytokine to induce pro-fibrotic associated changes. To better explore the bioenergetics status of these cells quantitatively, the oxygen consumption rate (OCR) and the extracellular acidification rate (ECAR) of HKC-8 cells were measured during sequential treatment with compounds that impact mitochondrial activity using a Seahorse XF24 Extracellular Flux Analyzer. After 48h of treatment with TGF-β1, HKC-8 cells showed a consistent decrease in OCR, at the different stages of cellular respiration in the presence of miR-NC. By contrast, over-expression of miR-9-5p was able to counteract the TGF-β1-induced reduction in mitochondrial respiratory capacity (Figure 7, A and B). When this same approach was used to verify the effect of miR-9-5p on glycolytic function by determining the ECAR, we found that miR-9-5p also prevented the inhibition of overall glycolytic function induced by TGF-β1 (Figure 7, C and D). Decreased glycolytic flux in TGF-β1-treated cells was reflected in a significant reduction (40%) in lactate production after 48h (Figure 7E). HKC-8 cells transfected with miR-9-5p showed almost identical amounts of lactate after TGF-β1 treatment than cells in basal conditions (Figure 7E). Decreased mitochondrial respiration and glycolytic function in TGF-β1-treated HKC-8 cells resulted in a 40% decrease in the intracellular ATP content, which was prevented by over-expression of miR-9-5p (Figure 7F). These results strongly support that the protective action of miR-9-5p is mediated by an enhancement of mitochondrial function, in consistence with the gene expression data described above.

**Figure 7.**
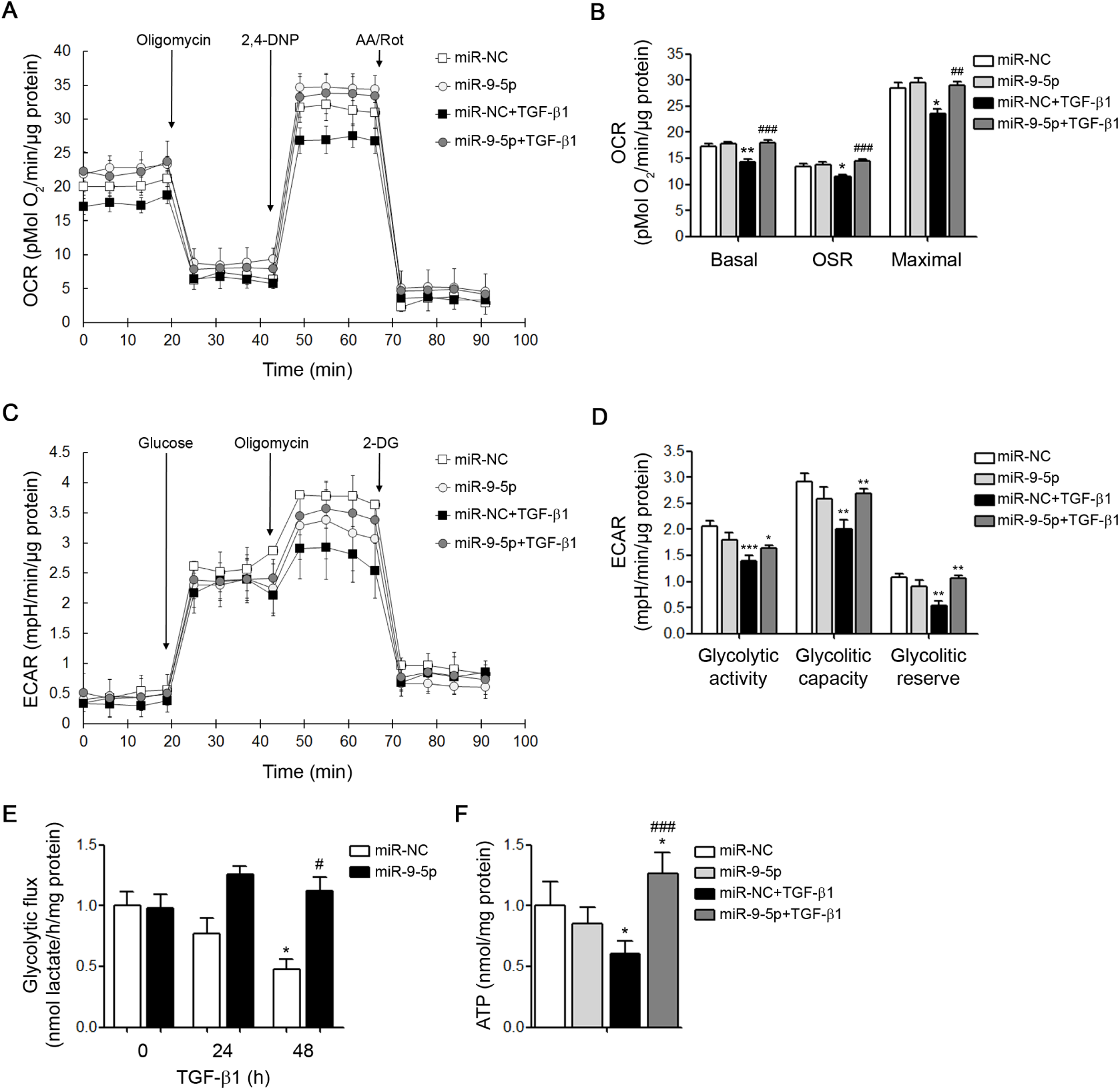
miR-9-5p prevents TGF-β1-induced metabolic and bioenergetics derangement in human kidney cells. (**A**) Human kidney tubular cells (HKC-8) were transfected with 40 nM miR-NC (control) or miR-9-5p and exposed to 10 ng/ml TGF-β1 for 48h. Oxygen consumption rate (OCR) was measured with a Seahorse XF24 Extracellular Flux Analyser. Oligomycin (6 µM), 2-4 dinitrophenol (2,4-DNP) (500 µM) and a combination of antimycin A (1 µM) and rotenone (1 µM) (AA/Rot) were injected sequentially at the time points indicated. (**B**) Bar graphs show the rates of basal respiration (Basal), oligomycin-sensitive respiration (OSR) and maximal respiration (Maximal) reflected by the OCR levels from (A). (**C**) Extracellular acidification rate (ECAR) of the HKC-8 cells treated as described in (**A**). Where indicated, glucose (10 mM), oligomycin (6 µM) and 2-deoxyglucose (2-DG, 100 mM) were added. (**D**) Bar graphs show the rates of glycolytic activity, glycolytic capacity and glycolytic reserve reflected by the ECAR levels. (**A** and **C**) Each data point represents the mean ± SEM of 4-6 independent experiments. (**B** and **D**) Data represent mean ± SEM from 4-6 independent experiments. (**A**-**D**) Data are represented after normalization by protein amount. (**E**) Glycolytic flux reflected by lactate production rate was measured in HKC-8 cell culture supernatants transfected with 40 nM miR-NC (control) or miR-9-5p and exposed to 10 ng/ml TGF-β1 for the indicated time points. Bar graphs represent mean ± SEM from 4 independent experiments. (**F**) Total cellular ATP concentration in HKC-8 cells treated as described in (**A**). Graph bars show mean ± SEM from 4-5 independent experiments. (**E** and **F**) Data were normalized by milligrams (mg) of protein. In all data, *P < 0.05, **P < 0.01, ***P < 0.001 compared to control cells; ^#^P < 0.05, ^##^P < 0.01, ^###^P < 0.001 compared to their corresponding negative control time point, non-parametric two-tailed Mann-Whitney U test.

### The protective role of miR-9-5p in kidney fibrosis is mediated by PGC-1α

The analysis of the RNA-Seq data with the DAVID gene annotation tool revealed that the PPAR signaling pathway was significantly affected by miR-9-5p in the injured kidney (Supplemental 4B, pink bar), suggesting the possible implication of this pathway in the protective role of miR-9-5p in kidney fibrosis. The complex formed by peroxisome proliferator activated receptor alpha (PPARα) and its co-activator PGC-1α controls the FAO gene program, having a direct impact on kidney fibrosis (49). We found that miR-9-5p was able to prevent the down-regulation of both *Ppargc1a* (Figure 8A) and *Ppara* (Figure 8B) genes 10 days after UUO. To probe the role of PGC-1α in the protective effect of miR-9-5p in renal fibrosis, null mice (PGC-1α ^−/−^) and their control counterparts (PGC-1α ^f/f^) were subjected to UUO for 7 days. MiR-9-5p prevented the up-regulation of the fibrotic expression markers *Acta2* and *Pdgfrb*, (Figure 8C) and of the ECM genes *Col1α*1 and *Fn1* (Figure 8D) in the obstructed kidneys of PGC-1α ^f/f^ mice. On the contrary, this effect was not observed in the PGC-1α ^−/−^ mice. Sirius red staining confirmed that the reduction in the collagen accumulation promoted by miR-9-5p was not significant in the PGC-1α ^−/−^ mice (Figure 8E). These results indicated that miR-9-5p requires PGC-1α to convey its protective role in renal fibrosis.

**Figure 8.**
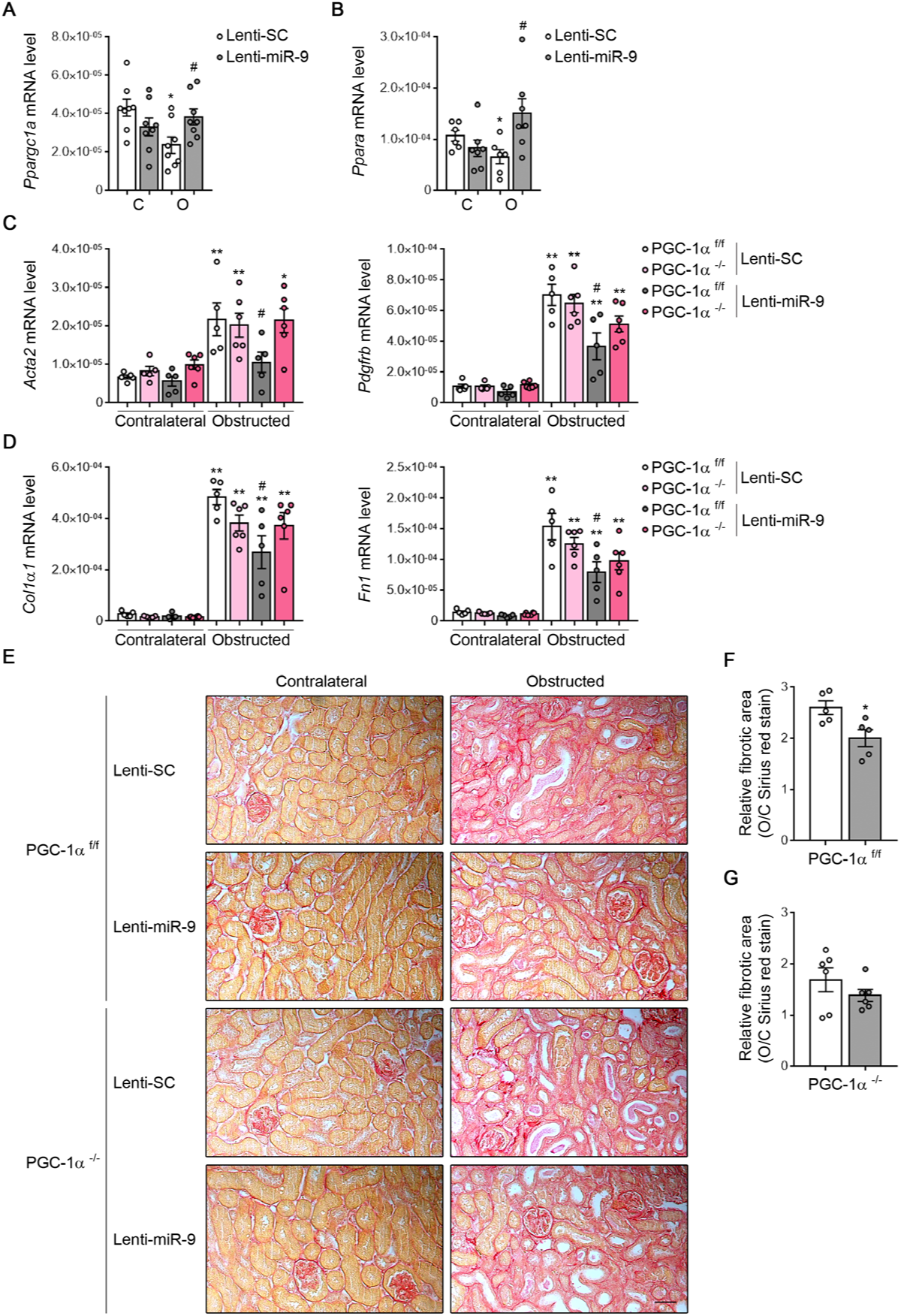
PGC-1α is necessary to sustain the anti-fibrotic effects of miR-9-5p. (**A** and **B**) Lentiviral particles (2×10^7^ ifu/mouse) containing miR-9-5p (Lenti-miR-9) or scramble miRNA (Lenti-SC) were injected to C57BL/6 mice and 5 days later they were subjected to UUO for 10 days. mRNA levels of peroxisome proliferator-activated receptor gamma coactivator 1 alpha (*Ppargc1a*) (**A**) and peroxisome proliferator-activated receptor alpha (*Ppara*) (**B**) genes in contralateral (C) and obstructed (O) kidneys from mice described above were analyzed by qRT-PCR. (**C** and **D**) PGC1-α ^−/−^ mice were injected as described above and 5 days later they were subjected to UUO for 7 days. PGC-1α ^f/f^ mice were used as controls. mRNA levels of *Acta2* (left panel), *Pdgfrb* (right panel) (**C**), *Col1α1* (left panel) and *Fn1* (right panel) (**D**) genes in kidneys from these mice were analyzed by qRT-PCR. (**A**-**D**) *P < 0.05, **P < 0.01 compared to their corresponding control kidneys; ^#^P < 0.05, compared to kidneys from mice administered Lenti-SC. (**E**) Representative microphotographs of one mouse per group of sirius red staining of kidneys from PGC1-α ^−/−^ and PGC1-α ^f/f^ treated as described in (**C** and **D**). Scale bar: 50 µm. (**F** and **G**) Quantification of sirius red staining was calculated as a ratio of the stained area over the total area. The fibrotic area is represented as the ratio of sirius red staining of obstructed (O) related to their corresponding contralateral (C) kidney (O/C) of mice treated as described above in. *P < 0.05, compared to O/C kidneys in mice administered Lenti-SC. (**A**-**D**, **F** and **G**) Statistical significance was determined using non-parametric two-tailed Mann-Whitney U test. Bar graphs show the mean ± SEM (n = 5-8 mice per group).

## Discussion

Kidney fibrosis remains a major challenge from clinical, therapeutic and biological standpoints. Clinically, it represents the final common pathway of most progressive nephropathies, with independence of their etiology. Therapeutically, in spite of major efforts and encouraging results based on angiotensin II antagonism (9, 50), no substantial advances have been made, while the possibility of reverting fibrosis seems distant. Biologically, besides important progress in the understanding of cellular and molecular mechanisms, several conundrums pervade the field. The discovery of miRNAs has fostered important research on their potential use as therapeutic agents or targets in almost every clinical setting, including kidney disease (33). In previous work (34, 35) we identified miR-9-5p as a miRNA with anti-fibrotic potential in the lung, peritoneum and skin. We now show that miR-9-5p exerts a protective effect in kidney fibrosis and provide data supporting a crucial role for metabolic reprogramming mediating this action (Figure 9).

**Figure 9.**
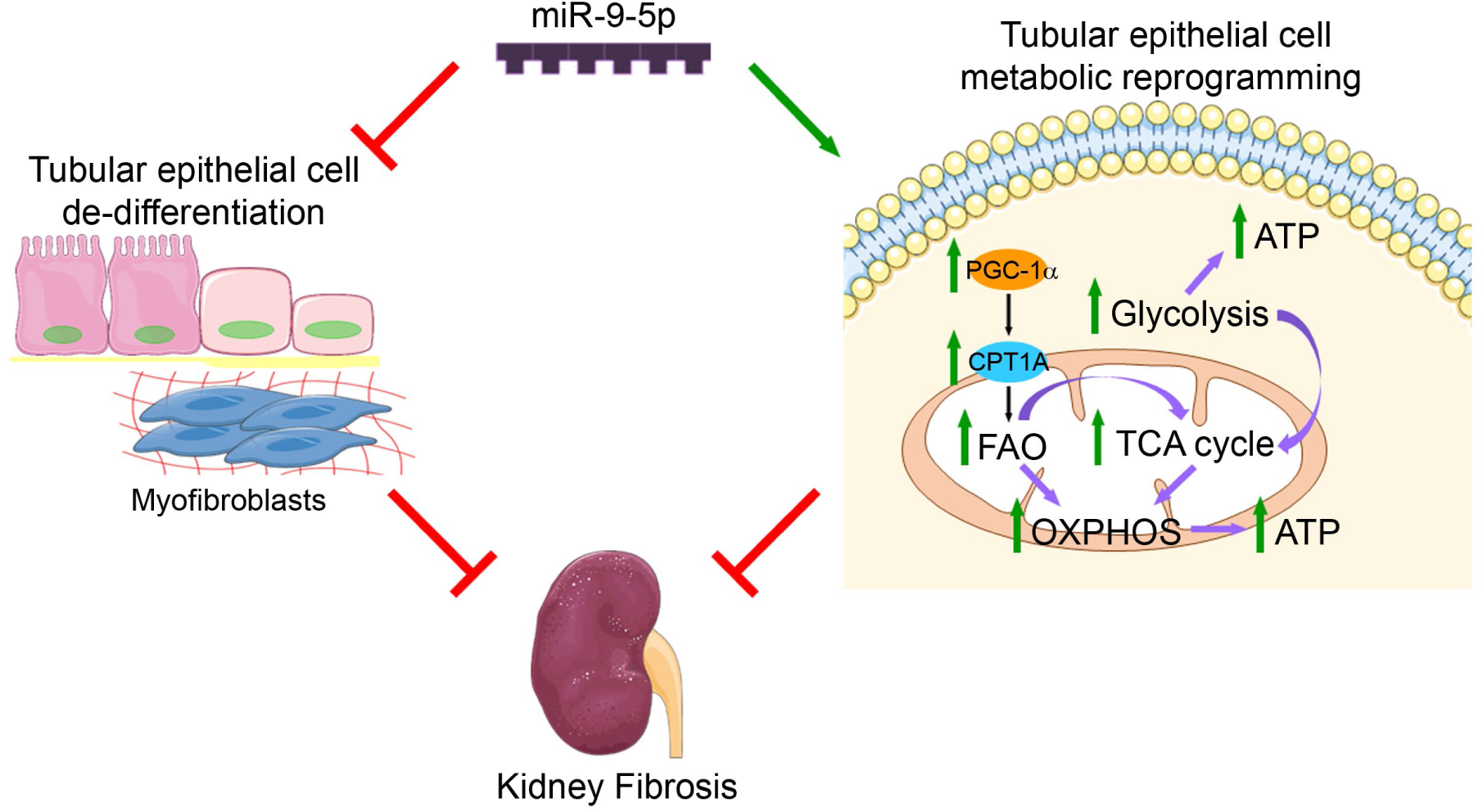
Protection of renal fibrosis promoted by miR-9-5p. Over-expression of miR-9-5p prevents tubular epithelial renal cell de-differentiation and reprograms fibrosis-related metabolic derangement.

Epithelial injury remains a central event in the pathogenesis of CKD. Direct epithelial damage causes de-differentiation and up-regulation of mediators that orchestrate pro-inflammatory signaling (51) and activate interstitial myofibroblasts in a paracrine fashion (8). We found that miR-9-5p not only mitigated the conspicuous histopathological changes associated to UUO, but also significantly diminished the presence of cellular markers associated to tubular cell transformation and inflammation. Moreover, miR-9-5p reduced epithelial cell injury and macrophage accumulation in UUO-associated fibrosis. Studies in HKC-8 confirmed that miR-9-5p exerts protection at the cellular level, most likely by blunting TGF-β1-dependent signaling pathway. Previously identified targets for miR-9-5p in the lung such as *Tgfbr2* and *Nox4* were also confirmed in the UUO model. As in the lung, kidney injury was also associated with an increment in miR-9-5p levels, likely as part of a homeostatic response towards counteracting TGF-β-mediated damage (34). These observations support a protective action of miR-9-5p on epithelial tubular cells by attenuating their de-differentiation.

Beyond the paramount relevance of TGF-β1 and other pro-fibrogenic factors as canonical pathways in kidney fibrosis, two important studies have emphasized the crucial role of FAO and mitochondrial function in renal fibrosis (3) (23). We found that miR-9-5p counteracted the down-regulation of quintessential metabolic pathways related to FAO and mitochondrial function promoted by UUO. In particular, the repression of genes related to both glycolysis and FAO elicited by UUO was clearly mitigated by miR-9-5p. Of importance, we observed that miR-9-5p prevented the down-regulation of CPT1A, a crucial enzyme for shuttling long chain fatty acids into the mitochondria. This attests to a specific capacity for miR-9-5p to regulate an ample metabolic response preserving energy availability in the kidney. Nevertheless, the precise mechanisms or targets by which miR-9-5p may alter the expression of genes and their cognate transcripts involved in the aforementioned pathways, remain to be established. Several other miRNAs, including miR-29 (52, 53), miR-30c (54), miR-200 (55, 56), miR-324 (57) and miR-433 (58, 59) have been shown to modify the fibrotic phenotype by operating through different targets and mechanisms. However, metabolic reprogramming was not an essential component of their mode of action.

Alterations in FAO and mitochondrial bioenergetics have proven to be decisive in kidney disease, both in AKI and CKD (60-62). A significant degree of the pro-fibrotic actions attributed to miR-21 in the kidney are related to profound and specific metabolic derangements closely associated to mitochondrial dysfunction (10). Up-regulation of genes related to mitochondrial function and to the PPARα signaling pathway was observed by anti-miR-21 treatment (32). By contrast, we found that miR-9-5p was able to exert opposite actions to those reported for miR-21 as miR-9-5p prevented TGF-β1-induced reductions in mitochondrial respiration, glycolytic function and ATP generation. Closely related to mitochondrial function and biogenesis is the critical role of the transcriptional co-activator PGC-1α, whose contribution to the preservation of kidney integrity in several models of damage is clearly established (49, 63). Tubular gain-of-function of PGC-1α protected mice against Notch-induced kidney fibrosis and reversed the mitochondrial derangements associated with this model (64). However, there are few reports addressing the effect of PGC-1α deficiency in chronic kidney damage. In our study, global deletion of PGC-1α was not associated with an exacerbated fibrotic phenotype. Nevertheless, the presence of PGC-1α seems indispensable to promote a miR-9-5p-dependent protective response, in consistence with the observed amelioration of mitochondrial function.

A search of human and mouse miRNA tissue databases (65, 66) did not find hits for miR-9-5p in the kidney. Although UUO induced a moderate increase in miR-9-5p levels, this is unlikely to effectively deter the progression of fibrosis. In addition, the low basal levels of this miRNA in the kidney could limit the possibility to achieve beneficial effects by promoting its endogenous manipulation. In contrast, we found that exogenous delivery of miR-9-5p, even achieving only a modest enhancement of its expression, can be exploited to prevent and defer renal fibrosis. This also calls to the necessity of identifying more selective and efficient modes of miRNA delivery into the kidney. Thus, miRNAs with a beneficial action should be object of active research, as they may constitute future valid therapeutic alternatives for CKD.

## Methods

### Cell culture and treatments

Human proximal tubule epithelial cell line (HKC-8) was kindly provided by Katalin Susztak (Philadelphia, Pennsylvania, USA) (3). Cells were grown in Dulbecco’s Modified Eagle Medium: Nutrient Mixture F-12 (DMEM/F-12) containing 10% (vol/vol) fetal bovine serum (FBS) (HyClone Laboratories, Logan, UT), 1x Insulin-Transferrin-Selenium (ITS) (Gibco, Rockville, MD), 0.5 µg/ml hydrocortisone (Sigma-Aldrich, St. Louis, MO) and 1% (vol/vol) penicillin/streptomycin (Gibco) at 37°C in 5% CO2. For treatment, cells were serum starved overnight and then incubated with 10 ng/ml recombinant human transforming growth factor-beta 1 (TGF-β1) (R&D Systems, Minneapolis, MN) for indicated times.

### Transfection of miRNA precursors

Cells at 60% confluence were transfected with 40 nM of mirVana™ miRNA mimic of miR-9-5p (miR-9-5p) (4464066, Ambion, Carlsbad, CA) using Lipofectamine 2000 (Invitrogen, Carlsbad, CA). In all experiments, an equal concentration of a non-targeting sequence mirVana™ miRNA mimic Negative Control #1 (miR-NC) (4464058, Ambion) was used.

### RNA isolation

Total RNA was isolated from cells and kidney tissue using the miRNeasy Mini Kit (Qiagen, Valencia, CA). RNA quantity and quality were determined at 260 nm by a Nanodrop spectrophotometer (Nanodrop Technologies, Wilmington, DE).

### Analysis of mRNA expression by RT-PCR

For syber (SYBR) green assays, reverse transcription (RT) was carried out with 1 µg of total RNA using the iScript^TM^ cDNA Synthesis Kit (Bio-Rad, Hercules, CA). Quantitative reverse transcriptase polymerase chain reaction (qRT-PCR) was carried out with the iQ™SYBR Green Supermix (Bio-Rad), using a 96-well Bio-Rad CFX96 RT-PCR System with a C1000 Thermal Cycler (Bio-Rad). A Cq value was obtained from each amplification curve using CFX96 Analysis Software provided by the manufacturer. GAPDH, for cell samples, and 18S, for kidney mice samples, genes were used for normalization purposes. Gene expression fold changes were normalized to values of HKC-8 control cells. The primer sequences used for mRNA quantification are listed in Supplemental Table 5. For Taq-Man® assays, total RNA was reverse transcribed at a concentration of 20 ng/µl with the High Capacity RNA to cDNA kit (Thermo Fischer Scientific) for 2h at 37°C. Gene expression profile of selected genes was carried out in a HT-7900 Fast Real time PCR System using specific primers and Taq-Man® probes (Thermo Fischer Scientific), see Supplemental Table 6 for the specific codes. Cq values were calculated for each specific gene using common settings in all samples and data were analyzed with the StatMiner® software (Integromics, S.L., Perkin Elmer, Waltham, MA). An average of UBC and 18S rRNA was used for normalization purposes. In both, SYBR and Taq-Man® assays, relative mRNA expression was determined using the 2 ^−ΔΔCt^ method (67).

### Quantification of miRNA expression

Quantification of miR-9-5p expression in kidney mice samples was performed using the miRCURY LNA RT Kit (Qiagen). Following RT, the cDNA template was amplified using microRNA-specific miRCURY LNA^TM^ primers for mature miR-9-5p (YP00204513, Qiagen). qRT-PCR was performed as described above. Cq values were normalized to the endogenous control 5S rRNA (YP00203906, Qiagen). Relative miRNA expression was determined using the 2 ^−ΔΔCt^ method (67).

### Western blotting and antibodies

Briefly, cells were washed with PBS, homogenized and lysed in 100 µl RIPA buffer containing 150 mM NaCl, 0.1% SDS, 1% sodium deoxycholate, 1% NP-40 and 25 mM Tris-HCl pH 7.6, in the presence of protease (Complete, Roche Diagnostics, Mannheim, Germany) and phosphatase inhibitors (Sigma-Aldrich). Cells were harvested by scraping and the samples were clarified by centrifugation at 13,000 rpm for 15 min at 4°C. For mouse sample protein extraction, kidneys were homogenized in the 300 µl RIPA buffer above described and disrupted by using the TissueLyser LT (Qiagen). The mixture was spun down (15 min at 14,000 rpm) at 4°C. The pellet was then discarded and the supernatant was kept as kidney lysate. Protein concentrations were determined by the BCA Protein Assay Kit (Thermo Fisher Scientific, Rockford, IL). Equal amounts of protein (10-50 µg) from the total extract were separated on 8-10% SDS-polyacrylamide gels and transferred onto nitrocellulose blotting membranes (GE Healthcare, Germany) at 12 V for 20 min in a semi-dry Trans-Blot Turbo system (Bio-Rad). Membranes were blocked by incubation for 1 hour with 5% non-fat milk in PBS containing 0.5% Tween-20, and blotted with specific antibodies to α-Actin (1A4) (1:1,000, sc-32251, Santa Cruz Biotechnology, Dallas, Texas), Fibronectin (1:1,000, F7387, Sigma-Aldrich), CPT1A (1:500, ab128568, Abcam, Cambridge, UK) and phosphorylated Smad2 (pSmad2, Ser465/Ser467) (1:1,000, 3101, Cell Signaling, Danvers, MA). After incubation with IRDye 800 goat anti-rabbit and IRDye 600 goat anti-mouse (1:15,000, LI-COR Biosciences, Lincoln, NE) secondary antibodies, membranes were imaged in triplicates with the Odyssey Infrared Imaging System (LI-COR Biosciences). Band sizing was performed using the ImageJ 1.5 software (http://rsb.info.nih.gov/ij) and relative protein expression was determined by normalizing to GAPDH (1:10,000, MAB374, Millipore, Bedford, MA), for cell samples, and to α-Tubulin (1:5,000, T9026, Sigma-Aldrich), for kidney mice samples. Phospho-Smad2 activity was calculated by normalizing to total Smad2 (D43B4) (1:1,000, 5339, Cell Signaling).

### Measurement of oxygen consumption and extracellular acidification rates

Oxygen consumption rate (OCR) and extracellular acidification rate (ECAR) in HKC-8 cells were studied by using a Seahorse XF24 Extracellular Flux Analyzer (Agilent, Santa Clara, CA). Key parameters of mitochondrial function were assessed by directly measuring the OCR in HKC-8 cells transfected with 40 nM of either miR-9-5p or miR-NC and treated with 10 ng/ml TGF-β1 for 48h. Then, cells were seeded in a XF24 V7 cell culture microplate (Agilent) at 1×10^4^ cells per well and equilibrated with Seahorse XF Base Medium (Agilent) supplemented with 1 mM pyruvate, 2 mM glutamine and 10 mM glucose for 1h in a CO2 free incubator previous to the assay. Mitochondrial function was determined through sequential addition of 6 μM oligomycin, 500 µM 2-4 dinitrophenol (2,4-DNP) and 1 μM antimycin/1 μM rotenone. This allowed to measure the basal respiration (basal), that shows the energetic demand of cells under baseline conditions; the Oligomycin Sensitive Respiration (OSR), calculated as the difference between the basal respiration and the OCR measured as described above, after the addition of oligomycin and represents the ATP-linked oxygen consumption and maximal respiration (maximal) that represents the maximal OCR attained after adding the uncoupler 2,4-DNP. Quantification of these values where done by normalizing from basal values and after correction for non-mitochondrial OCR. Glycolytic function was determined by measuring the ECAR. HKC-8 cells were treated and seeded as indicated above. Cells were equilibrated with Seahorse XF Base Medium (Agilent) supplemented with 1 mM glutamine for 1h previous to extracellular flux assay. Glycolitic function was determined through sequential addition of 10 mM glucose, 6 µM olygomycin and 100 mM 2-deoxyglucose (2-DG). This allowed to assess key parameters of glycolytic flux as glycolysis, measured as the ECAR reached by cells after the addition of saturating amounts of glucose; glycolytic capacity, that is the maximum ECAR reached after addition of oligomycin, thus shutting down oxidative phosphorylation and driving cells to use glycolysis to its maximum capacity and glycolytic reserve, that indicates the capability of cells to respond to an energetic demand as well as how close the glycolytic function is to the celĺs theoretical maximum. Quantification of these values where done by normalizing from basal values and after correction for non-glycolytic acidification. All injected compounds were prepared in the same medium in which the experiment was conducted and were injected from the reagent ports automatically at indicated times. OCR and ECAR data were normalized to total cellular protein.

### Lactate Production measurement

The initial rates of lactate production were determined as an index of aerobic glycolysis by enzymatic determination of lactate concentrations in the culture medium of HKC-8 cells transfected with 40 nM of either miR-9-5p or miR-NC and treated with 10 ng/ml TGF-β1 for 24 and 48h. Culture medium was replaced by fresh medium 1h before the measurement. Samples were prepared by the addition of 800 µl cold perchloric acid (Merck, Darmstadt, Germany), to stop lactate dehydrogenase (LDH) enzymatic activity, to 200 µl of culture medium, incubated on ice for 1h and then centrifuged for 5 min, 11,000 g at 4°C to obtain a protein-free supernatant. The supernatants were kept on ice and neutralized with 20% (w/v) KOH (Merck) and centrifuged at 11,000 g and 4°C for 5 min to sediment the KClO4 salt. Lactate levels in the clear supernatants were determined using a LDH-based enzymatic assay (68). The rate of glycolytic flux was expressed in nmol lactate/h/mg protein.

### ATP measurement

Cellular ATP concentrations were determined using the ATP Bioluminescence Assay Kit CLS II (11 699 709 001, Roche) following the manufacturer’s instructions, on HKC-8 cells transfected with 40 nM of either miR-9-5p or miR-NC and treated with 10 ng/ml TGF-β1 for 48h. The rate of ATP cellular production was expressed in nmol ATP/mg protein.

### Lentiviral vector construct

The Lentivector-based miRNA precursor constructs expressing the miR-9-5p (Lenti-miR-9) (MMIR-9-1-PA-1) and the scramble negative control (Lenti-SC) (MMIR-000-PA-1) were purchased from SBI System Biosciences (Mountain View, California). Pseudoviral particles were prepared using the pPACK-F1 Lentivector packaging system (SBI System Biosciences) and HEK293T producer cell line. The titration of pseudoviral particles generated with the lentiviral vectors was determined by determining the % of GFP^+^ cells by flow cytometry in a BD FACSCanto™ II High Throughput Sampler flow cytometer (Becton Dickinson Bioscience) 72h after infection. The lentiviral titers were calculated as described (69) and expressed in infection units (ifu)/ml.

### Animals

Males C57BL/6JRccHsd (C57BL/6) mice were purchased from Envigo (Horst, Netherlands). The generation and phenotype of C57BL/6J PGC-1α null mice (PGC-1α ^−/−^) mice were described previously (70). PGC-1α ^−/−^ mice were originally kindly provided by Dr. Bruce Spiegelman (Dana-Farber Cancer Institute, Harvard Medical School, Boston, MA, USA) and, following embryo transfer, a colony was established at the Instituto de Investigaciones Biomédicas “Alberto Sols” (IIB, Madrid, Spain) animal facility. C57BL/6J PGC-1α flox mice (PGC-1α ^f/f^), that were used as controls for the PGC-1α ^−/−^ mice experiments, were purchased from The Jackson Laboratory (https://www.jax.org/strain/009666, Bar Harbor, ME) and maintained in the IIB.

### Unilateral ureteral obstruction

The model of unilateral ureteral obstruction (UUO) was performed in 4-6-week-old males C57BL/6, PGC-1α ^−/−^ and PGC-1α ^f/f^ mice, under isoflurane-induced anesthesia. UUO was performed as described (71). Briefly, the left ureter was ligated with silk (5/0) at two locations and cut between ligatures to prevent urinary tract infection (obstructed kidney). miR-9-5p expression was evaluated in kidneys after UUO. To determine the effect of miR-9-5p over-expression in the UUO, 2×10^7^ ifu/mouse of Lenti-miR-9 diluted in 30 µl saline serum were retro-orbital injected in mice 5 days before UUO. As controls, mice were provided with the same volume of Lenti-SC. To evaluate the persistence of lentiviral transduction in our model, miR-9-5p expression level was determined by qRT–PCR. Mice were subjected to surgery 5 days after lentiviral particles injection, they were sacrificed at the indicated time points and the PBS-perfused kidneys were harvested for analysis. At the time of sacrifice, animals were anesthetized with 5 mg/kg xylazine (Rompun, Bayer AG, Leverkusen, Germany) and 35 mg/kg ketamine (Ketolar, Pfizer, New York, NY). The left (obstructed) and the right (contralateral) kidneys were compared in each mouse in all experiments. A control group of sham-operated mice was assessed, showing similar results as contralateral kidneys in all the analysis performed (data not shown).

### RNA sequencing

To determine the global gene-expression and potential mechanisms involved in the miR-9-5p-induced changes during UUO, RNA sequencing (RNA-Seq) analysis was performed in kidneys from C57BL/6 mice retro-orbitally injected with 2×10^7^ ifu/mouse of Lenti-miR-9 diluted in 30 µl saline serum 5 days before UUO (Group 3: Contralateral and Group 4: Obstructed kidneys). As controls, C57BL/6 mice were provided with the same volume of Lenti-SC (Group 1: Contralateral and Group 2: Obstructed kidneys). 5 days after UUO, RNA was extracted and treated with DNase I (New England Biolabs, Ipswich, MA) to remove any possible contaminant of DNA. Libraries were prepared according to the instructions of the Kit “NEBNext Ultra Directional RNA Library Prep kit for Illumina” (E7760L, New England Biolabs, Ipswich, MA), following the protocol “Poly(A) mRNA Magnetic Isolation Module” (New England Biolabs). The input yield of total RNA to start the protocol was 1 µg, estimated by Agilent 2100 Bioanalyzer using a RNA 6000 nano LabChip kit (Agilent). The fragmentation time used for RNA samples was 8-10 min. We performed the library amplification included in the mentioned protocol using a PCR of 13 cycles. The obtained libraries were validated and quantified by an Agilent 2100 Bioanalyzer using a DNA7500 LabChip kit (Agilent) and an equimolecular pool of libraries was prepared and titrated by qRT-PCR using the “Kapa-SYBR FAST qPCR kit forLightCycler480” (Kapa BioSystems, Fritz Hoffmann-La Roche, Basilea, Switzerland) and a reference standard for quantification (Genomics Unit, Science Park, Madrid). The pool of libraries was denatured prior to be seeded on a NextSeq flowcell (Illumina, San Diego, CA), at a density of 2,2 pM, where clusters were formed and sequenced using a “NextSeq™ 500 High Output Kit” (Illumina), in a 1×75 single read sequencing run. An average of 32.5 million of pass-filter reads were obtained (range 28.4 – 36.0 · 106) and used for further filtering and bioinformatics analysis. Sequences were aligned against Mm version9 genome using TopHat aligner (G_PRO 2.0, Biotechvana, Valencia, Spain) and gene expression profiles, calculated in terms of fragments per kilobase of exon model per million reads mapped (FPKM) values, were determined using CuffDiff2. The sequences were deposited in the European Nucleotide Archive (ENA, https://www.ebi.ac.uk/ena) under the study accession number PRJEB32662 with the ID of the reads from ERS3424621 (SAMEA5620536) to ERS3424632 (SAMEA5620547).

### RNA-Seq data analysis

Genes which were detected in very low amounts (filtered out when total counts within the set of samples was lower than 120 counts and when the value of the relative counts were 0 in any of the performed comparisons) were removed from the analysis. A total set of 13,232 genes was selected for profiling. The number of significant genes (Q-value ˂ 0.01, corrected for multiple hypothesis testing with the Benjamini–Hochberg FDR algorithm (72)) in all the compared groups are showed in Supplemental Figure 4A. These genes were selected for further analysis by using the Venn’s diagrams drawing tool software Venny 2.1 (http://bioinfogp.cnb.csic.es/tools/venny, designed by Juan Carlos Olivero, BioinfoGP, CNB-CSIC) in order to graphically visualize the sets of genes resulting from all the possible compared groups. Genes resulted after this analysis are listed in Supplemental Table 1. The 949 genes resulted from the intersection between 2 vs 1 and 4 vs 2 were selected to perform Genes Functional Annotation Clustering in the bioinformatics Database for Annotation, Visualization and Integrated Discovery (DAVID) version 6.8 (http://david.abcc.ncifcrf.gov/) (73) using the Kyoto Encyclopedia of Genes and Genomes (KEGG) pathway databases. Gene Ontology annotation enrichment analysis, using the DAVID Bioinformatics Resources, was carried out to identify biological processes (BPs) represented by these changes in gene expression relative, thus making possible the enrichment of functionally related gene groups. Only BPs containing more than five genes involved in the term were considered. To gain insight into the enriched pathways regulated by miR-9-5p in the UUO, the same 949 genes described above were analyzed using the iPathwayGuide software (https://advaitabio.com/ipathwayguide/, AdvaitaBio, Plymouth, MI). Ingenuity Pathway Analyses (IPA, (https://www.qiagenbioinformatics.com/products/ingenuity-pathway-analysis/, IPA, Qiagen) was used for further analysis of the five metabolic pathways highlighted in the analysis described above included in the Ingenuity canonical pathways library. The significance of the association between the data set and the canonical pathway was measured in two ways, a percentage of the number of molecules from the data set that map to the canonical pathway and Fisher’s exact test used to calculate a P-value determining the probability that the association between the genes in the dataset and the canonical pathway. The stacked bar chart reveals the percentage of up- and down-regulated genes within each canonical pathway. In all data analysis, the enrichment score was reported as the minus log transformation of the geometric mean of P-values (modified Fisher’s exact test). The P-value was corrected for multiple hypothesis testing with the Benjamini–Hochberg FDR algorithm (72). Terms with a corrected P < 0.05 were considered significant.

### Histological and immunohistochemical analysis of mice kidney tissue

Kidney mice samples fixed in 4% (w/v) formaldehyde (415693, Carlo Erba reagents, Barcelona, Spain) were embedded in paraffin and cut into 3-5 μm tissue sections for immunohistochemistry (IHC) and histological studies. Paraffin-embedded kidney sections were stained using standard histology procedures (Hematoxylin and eosin (H&E), periodic acid-schiff (PAS) and sirius red). Immunostaining was carried out in 3 μm thick tissue sections. Sections were deparaffinized and exposed to the PT Link (Dako, Santa Clara, CA) with 10 mM citrate sodium buffer, pH 6.0-9.0 (depending of the immunohistochemical marker) for antigen retrieval. Samples were treated with 3% H2O2 to block endogenous peroxidase. Tissue sections were incubated with 4% bovine serum albumin (BSA) and 8% serum in 1x wash buffer ‘en vision’ (Dako) to eliminate non-specific protein binding sites. Primary antibodies, PCNA (1:500, sc-7907, Santa Cruz Biotechnology), Type I collagen (1:200, AB765P, Merck Millipore, Burlington, Massachusetts), α-Actin (1A4) (1:500, sc-32251, Santa Cruz Biotechnology) and F4/80 (1:100, MA1-91124, Thermo Fisher Scientific) followed by Immpress Reagent Kit (MP 7444, Vector, USA), were incubated overnight at 4°C. A biotinylated goat anti-rabbit IgG was applied to detect rabbit primary antibodies and complexes were visualized using the R.T.U Vectastain Elite ABC Kit (Vector Laboratories, Burlingame, CA). Mouse primary antibodies were visualized applying the VECTASTAIN® ABC HRP Kit (Peroxidase, Standard, Vector Laboratories, Burlingame, California). Tissue sections were revealed using 3,3’-diaminobenzidine (20 μl/ml, DAB, Dako) as chromogen and finally counterstained with Carazzi’s hematoxylin. Negative controls included non-specific immunoglobulin and no primary antibody. Finally, slides were mounted with mowiol (Merck Millipore). Images of H&E, PAS and sirius red were taken with bright field at 20x magnification using the KS300 imaging system, version 3.0 (Zeiss, Oberkochen, Germany) and the Axioskop2 plus microscope (Zeiss) with the DMC6200 imaging system (Leica Microsystems, Wetzlar, Germany). Histopathological analysis was performed by PAS staining in mice paraffin-embedded kidney samples in a blinded fashion by a pathologist. The tubular atrophy and tubular dilation was evaluated as previously described (50). The collagen deposition was evaluated by sirius red staining. The intensity of sirius red as well as the PCNA, α-SMA and F4/80 staining area was assessed with Image-Pro Plus 6.0 software (Media Cybernetics, http://www.mediacy.com/imageproplus). For each sample, processed in a blinded manner, the average value was obtained from the analysis of 3-6 non-overlapping fields (20x magnification) as density/mm2 of stained area as a percentage of the total analyzed area. Data of sirius red and F4/80 are expressed as the stained area of the obstructed kidneys compared to its corresponding contralateral kidneys.

### Flow cytometry analysis

Single cell suspensions were prepared from mice kidneys. Tissues were minced with a blade and digested with 100 mg/ml collagenase A and 10 μg/ml DNase I in PBS for 1h at 37°C, with agitation. After digestion, samples were passing through a 40 µm cell strainer and centrifuged 10 min, 2500 rpm at 4°C. The supernatants were removed and the pellets were resuspended in 500 µl fluorescence-activated cell sorting (FACS) buffer containing 1% FBS and 0.5% BSA. The antibodies used were: APC anti-human CD45 (Clone 2D1, 368511), APC/Cyanine7 anti-mouse F4/80 (Clone BM8, 123117), APC/Cy7 anti-mouse CD326 (Ep-CAM) (Clone G8.8, 118217), PE anti-mouse CD24 (Clone M1/69, 101807), all from BioLegend (San Diego, CA). Fc receptors were blocked with CD16/CD32 (clone 2.4G2, BD Biosciences). Cells (1×10^7^) were incubated with each antibody in FACS buffer in darkness, at room temperature for 20 min. Cells were then washed twice with FACS buffer and immediately analyzed on a FACSCanto flow cytometer with DIVA software (BD Biosciences). Dead cells were eliminated from analysis using Dapi (268298, Merck). For each experiment, we performed flow minus one (FMO) controls for each fluorophore to establish gates (APC Rat IgG2b, κ Isotype Ctrl Antibody, Clone RTK4530, 400611; APC/Cy7 Rat IgG2a, κ Isotype Ctrl Antibody, Clone RTK2758, 400523; PE Rat IgG2b, κ Isotype Ctrl Antibody, Clone RTK4530, 400607; all from Biolegend). Data analysis was performed using FlowJo v 10.0.8 software (https://www.flowjo.com, FlowJo, LLC, Ashland, OR).

### Statistical analysis

Differences between only two groups were analyzed statistically with non-parametric two-tailed Mann-Whitney U test or with the non-parametric Wilcoxon matched pairs signed rank test, when the control cells were normalized to 1. Data were analyzed with use of version 7.00 of the GraphPad Prism package (La Jolla, CA). A value of P < 0.05 was considered to be statistically significant (*P < 0.05, **P < 0.01, ***P < 0.01, ^#^P < 0.05, ^##^P < 0.01, ^###^P < 0.01). Data are reported as mean ± standard error of the mean (SEM).

### Study approval

Animals were handled in agreement with the Guide for the Care and Use of Laboratory Animals contained in Directive 2010/63/EU of the European Parliament. Approval was granted by the local ethics review board of Centro de Biología Molecular “Severo Ochoa” and of Instituto de Investigaciones Biomédicas “Alberto Sols”, after complying with the legal and ethical requirements relevant to the procedures for animal experimentation established by the Comunidad de Madrid.

## Author contributions

SL conceived and directed research. MFF designed, performed and analyzed the majority of experiments. VM, LME and CNT performed experiments. JIH, EBR and JT provided technical assistance for mouse experiments. CC and PC performed histological evaluation. MM provided the PGC-1α null mice. MRO provided intellectual and technical insight. RR performed the RNA-Seq and helped with the bioinformatics analysis. All authors helped with the discussion of the results and SL and MFF wrote the manuscript.

## Acknowledgements

This work was supported by Grants from the Ministerio de Economía y Competitividad (MINECO) SAF 2012-31388 (SL) and SAF2015-66107-R (SL), both cofunded by the European Regional Development Fund, Instituto de Salud Carlos III REDinREN RD12/0021/0009 and RD16/0009/0016 (SL and MRO), PI17/00119 (MRO), SAF2015-63904-R (MM), cofunded by the European Regional Development Fund, European Union’s Horizon 2020 research and innovation programme under the Marie Skłodowska-Curie grant agreement 721236-TREATMENT (MM), Comunidad de Madrid “NOVELREN” B2017/BMD-3751 (SL and MRO), and Fundación Renal “Iñigo Alvarez de Toledo” (SL), all from Spain. The CBMSO receives institutional support from Fundación “Ramón Areces”. VM and CNT were supported by pre-doctoral fellowships of the FPI Program (BES-2013-065986 and BES-2014-068929) from MINECO. We are grateful to the laboratories of Jorgina Satrustegui, especially to Araceli del Arco, and José Manuel Cuezva, for sharing the Seahorse equipment and their technical assistance and to Javier Casares from the laboratory of Miguel Ángel Alonso for his help in microscope image quantification, all from CBMSO. We are also grateful to Raúl Rodrigues Díez for his help in bioinformatics analyses and Laura Santos for her help with histology procedures, from the laboratory of Marta Ruiz-Ortega at the Fundación Jiménez Díaz. We deeply thank the work and collaborative spirit of the following facilities of the CBMSO: animal housing, flow cytometry, confocal microscopy, cell culture, ICT and the genomics and NGS core facility (CEI UAM+CSIC) that helped in the analysis with IPA software provided the data submission to a public repository. MFF and SL filed a patent in 2014 for the therapeutic use of miR-9-5p in fibrosis, which is not currently under commercial exploitation.

## Supplemental Figures and Supplemental figure legends

**Supplemental Figure 1.**
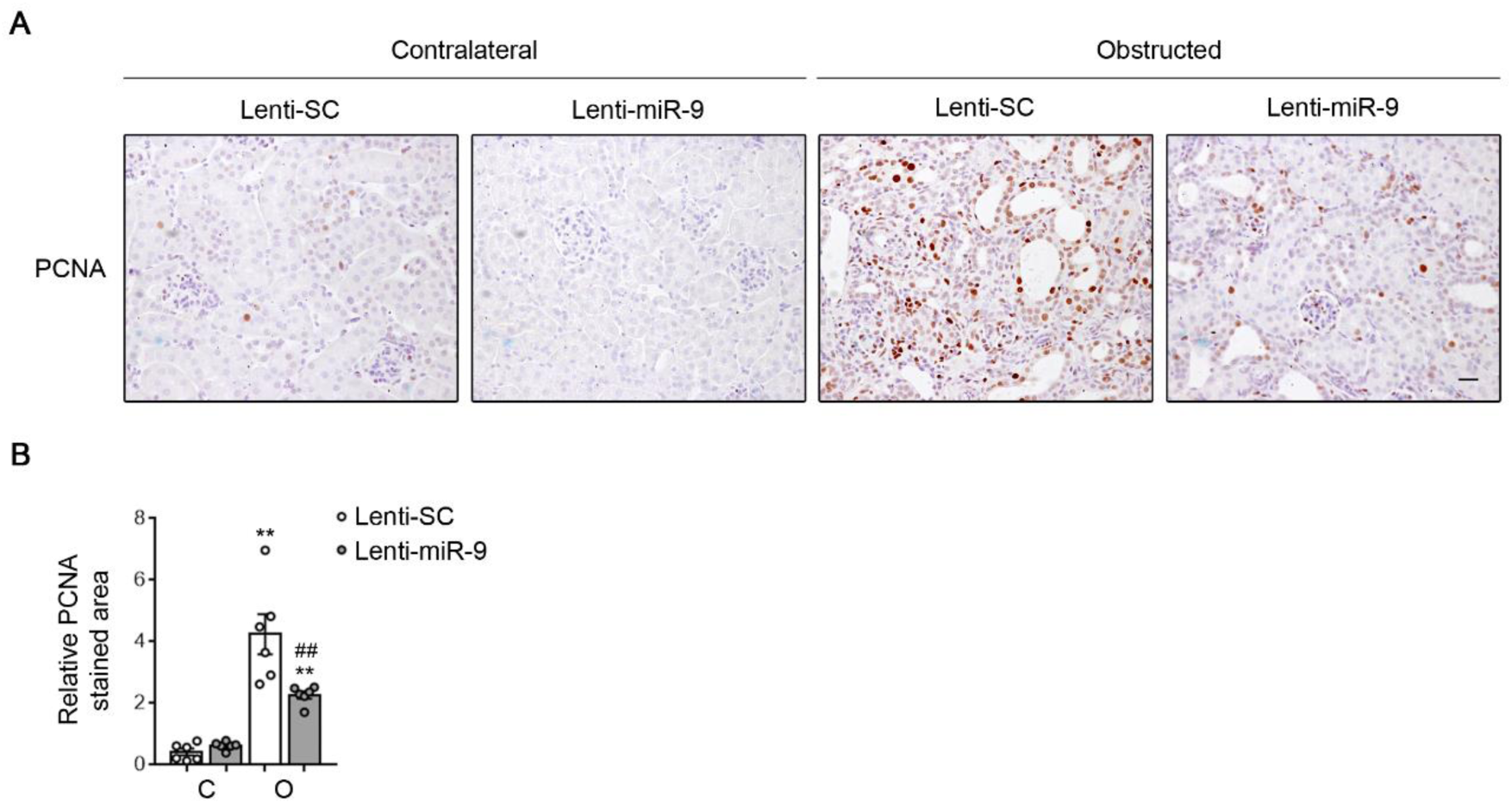
Over-expression of miR-9-5p decreases cell proliferation in UUO kidneys. (**A**) Representative microphotographs of one mouse per group of immunohistochemical staining for the Proliferating Cell Nuclear Antigen (PCNA) from kidney sections of C57BL/6 mice injected with lentiviral particles (2×10^7^ ifu/mouse) carrying miR-9-5p (Lenti-miR-9) or scramble miRNA (Lenti-SC) for 5 days. Kidneys samples were analyzed 10 days after UUO. Microscopic images are representative of at least 4 mice per group. Scale bar: 50 µm. (**B**) Bar graphs represent the quantification of PCNA positive stained area relative to the total analyzed area in the contralateral (C) and the obstructed kidneys (O) from mice treated as described in (**A**). Data represent the mean ± SEM. **P < 0.01 compared to their corresponding contralateral kidneys; ^##^P < 0.01 compared to obstructed kidneys in mice administered Lenti-SC, non-parametric two-tailed Mann-Whitney U test.

**Supplemental Figure 2.**
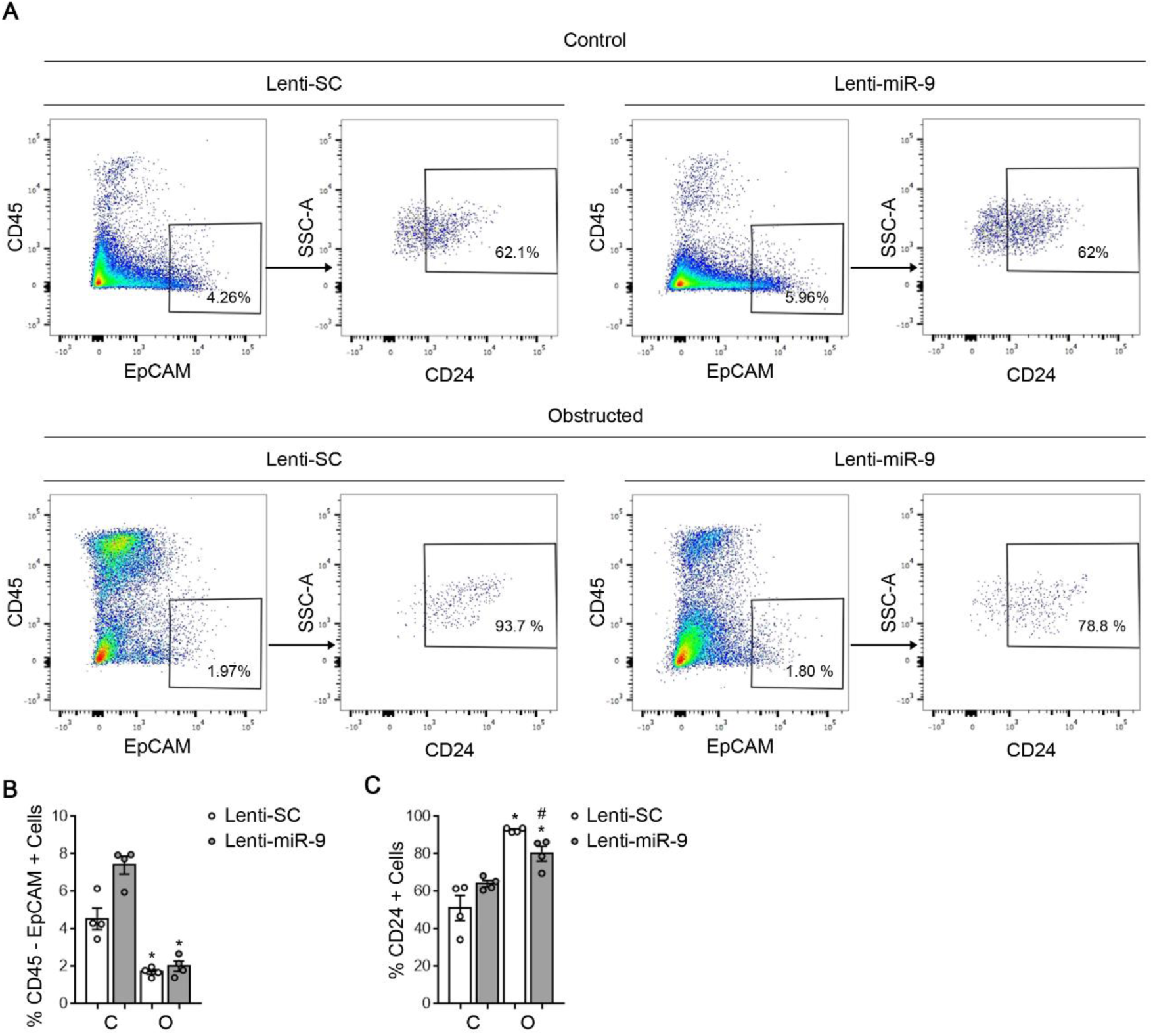
Administration of miR-9-5p reduces de-differentiated epithelial cell population in kidney fibrosis. (**A**) Examples of multiparameter flow cytometry dot plots illustrating cell population from contralateral (C) and obstructed (O) kidneys in mice subjected to UUO. Cells were gated for CD45^−^/Epithelial Cell Adhesion Molecule (EpCAM)^+^ and then selected for the presence of CD24. Cells analyzed correspond to disrupted tissue from mice injected with lentivirus (2×10^7^ ifu/mouse) containing miR-9-5p (Lenti-miR-9) or scramble miRNA (Lenti-SC) for 5 days and then subjected to UUO for 5 days. Numbers in quadrants indicate cell proportions in percent. SSC-A: Side Scatter Area. Data shown are representative of 4 mice per group. (**B**). (**B** and **C**) Data represent the mean ± SEM (n = 4 mice per group). *P < 0.05 compared to their corresponding contralateral kidneys; ^#^P < 0.05 compared to obstructed kidneys in mice administered Lenti-SC, non-parametric two-tailed Mann-Whitney U test.

**Supplemental Figure 3.**
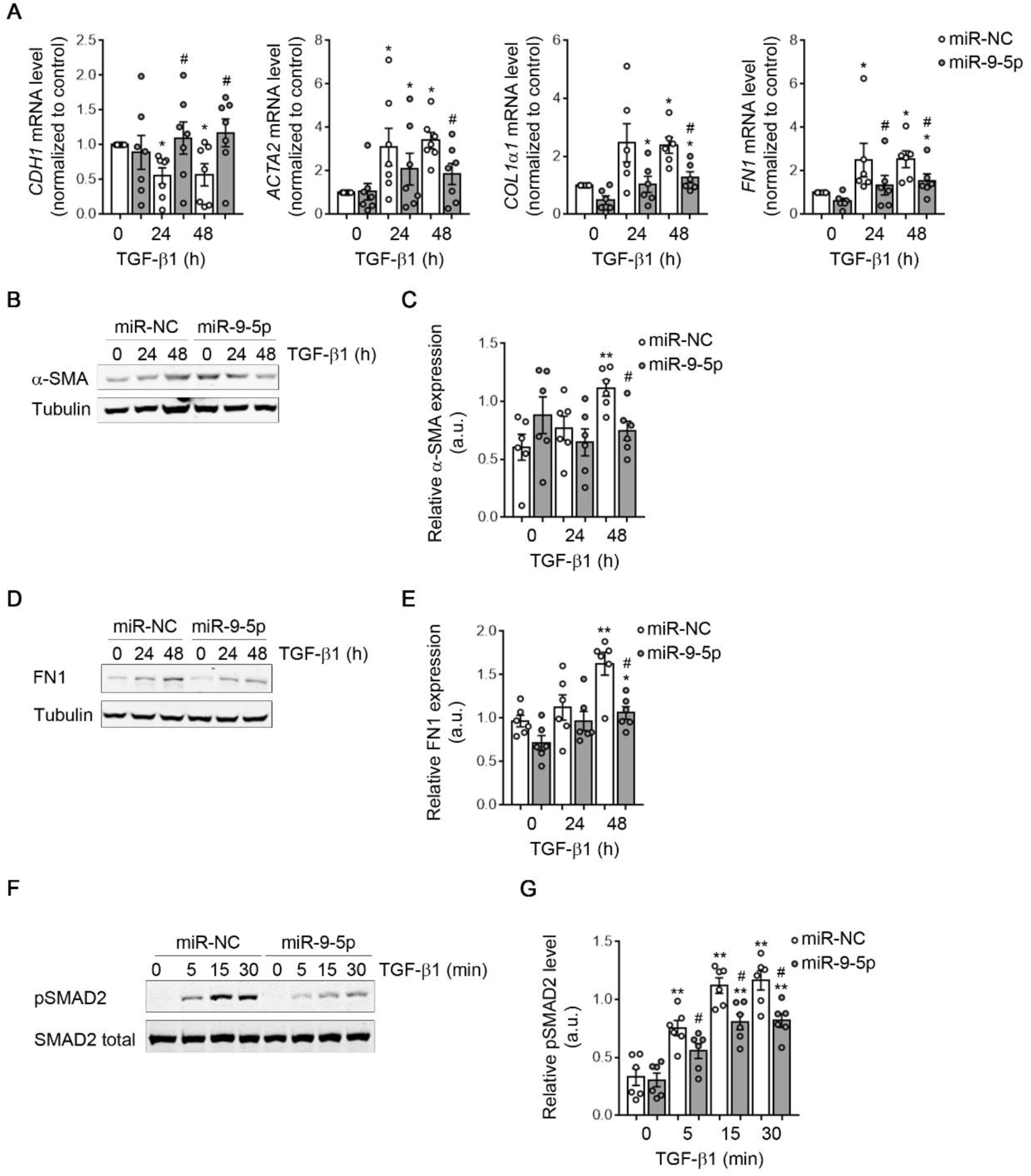
miR-9-5p administration results in delayed TGF-β1-induced fibrogenic transformation and Smad2 phosphorylation in human tubular epithelial cells. (**A**) mRNA expression of *CDH1*, *ACTA2*, *COL1α1* and *FN1* fibrotic markers were determined by qRT-PCR in HKC-8 cells transfected with 40 nM miR-NC (control) or miR-9-5p and exposed to 10 ng/ml TGF-β1 for the indicated time points. Data are represented after normalization to values of miR-NC-transfected cells. (**B** and **D**) Representative western blots of α-SMA and FN1 protein levels in HKC-8 cells treated as described in (**A**). (**C** and **E**) Bar graphs represent densitometric values (arbitrary units, a.u.) of protein expression. (**B**-**E**) Tubulin was used for normalization purposes. (**F**) Representative western blot of pSMAD2 protein levels in HKC-8 cells transfected as described in (**A**) and exposed to 10 ng/ml TGF-β1 for the indicated time points. (**G**) Bar graphs show densitometric values (arbitrary units, a.u.) of SMAD2 phosphorylation. (**F** and **G**) Total levels of SMAD2 were used for normalization purposes. (**A**, **C**, **E** and **G**) Data represent the mean ± SEM of 6-7 independent experiments. *P < 0.05, **P < 0.01 compared to cells transfected at time 0; ^#^P < 0.05 compared to their corresponding negative control time point, (**A**) Wilcoxon matched pairs signed rank test, (**C**, **E** and **G**) non-parametric two-tailed Mann-Whitney U test.

**Supplemental Figure 4.**
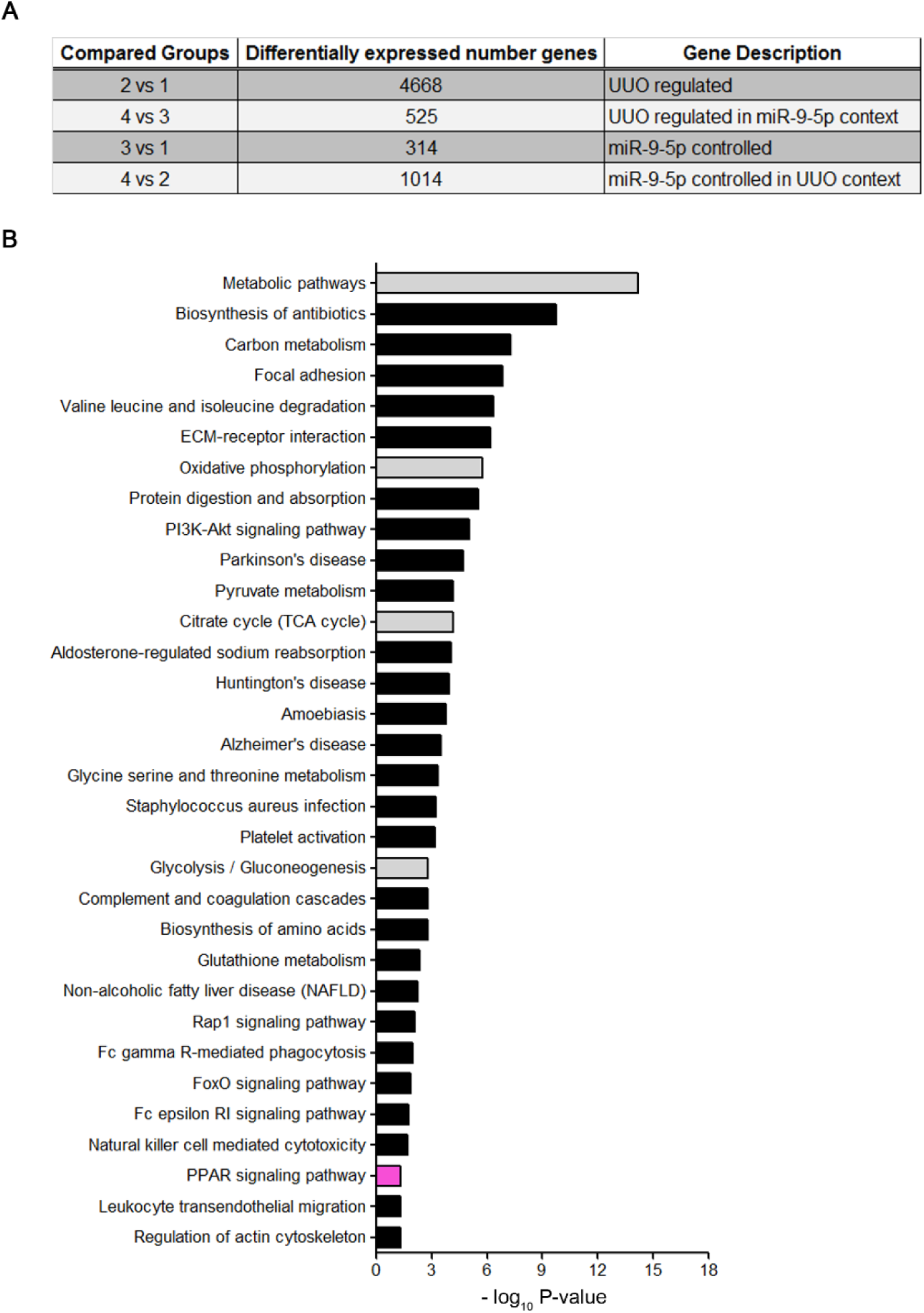
Cellular pathways involved in the differential gene expression related to UUO and miR-9-5p treatment. Lentiviral particles (2×10^7^ ifu/mouse) containing miR-9-5p (Lenti-miR-9) or scramble miRNA (Lenti-SC) were injected to mice and 5 days later they were subjected to UUO for 5 days. (**A**) The table shows the number of significant genes (Q-value ˂ 0.01) resulting from the comparisons among kidneys from mice groups treated as described above and as showed in Figure 5A. (**B**) KEGG enriched analysis showing the most prevalent biological pathways modified by the presence of miR-9-5p in mouse kidneys analyzed 5 days after UUO by using the DAVID database 6.8 software for analysis. The graph shows – log P-value calculated using modified Fisher’s exact test and corrected for multiple hypotheses testing with Benjamini-Hochberg FDR. Pathways with corrected P < 0.05 are depicted. Gray bars highlight metabolic related pathways and pink bar represents the PPAR signaling pathway.

**Supplemental Figure 5.**
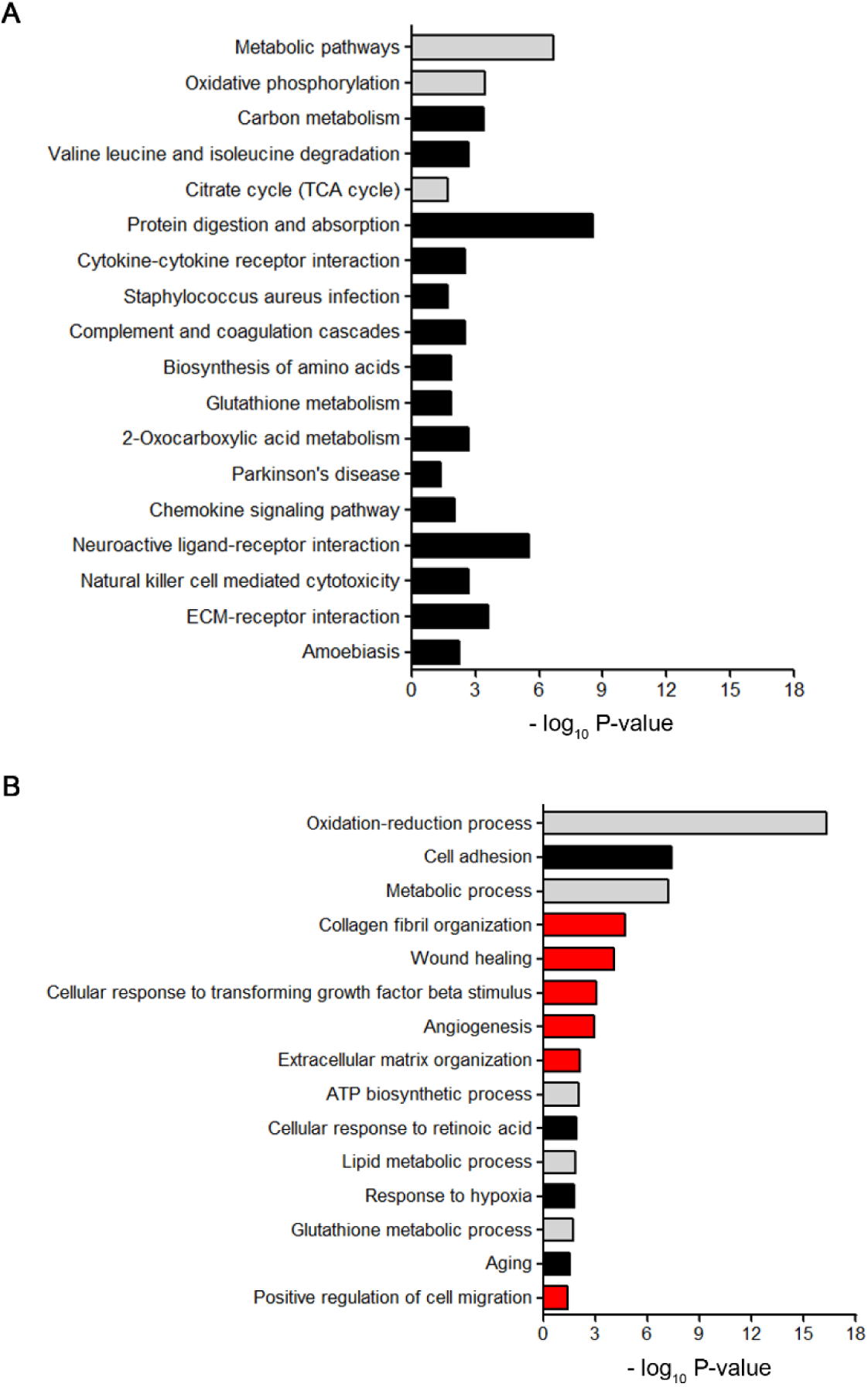
Main pathways and biological processes (BPs) modified by miR-9-5p in fibrotic kidneys. Lentiviral particles (2×10^7^ ifu/mouse) containing miR-9-5p (Lenti-miR-9) or scramble miRNA (Lenti-SC) were injected to mice and 5 days later they were subjected to UUO for 5 days. (**A**) KEGG enriched analysis showing the most prevalent biological pathways modified by the presence of miR-9-5p in kidneys from mice subjected to UUO for 5 days by using the using the iPathwayGuide software for analysis. Gray bars highlight metabolic related pathways. (**B**) Gene Ontology annotation enrichment analysis, using the DAVID database 6.8, was carried out to identify biological processes as indicated. Gray and red bars highlight metabolic- and fibrogenic-related processes respectively. (**A** and **B**) P-values were determined with modified Fisher’s exact test and corrected for multiple hypotheses testing with Benjamini-Hochberg FDR. Pathways with corrected P < 0.05 are depicted.

**Supplemental Figure 6.**
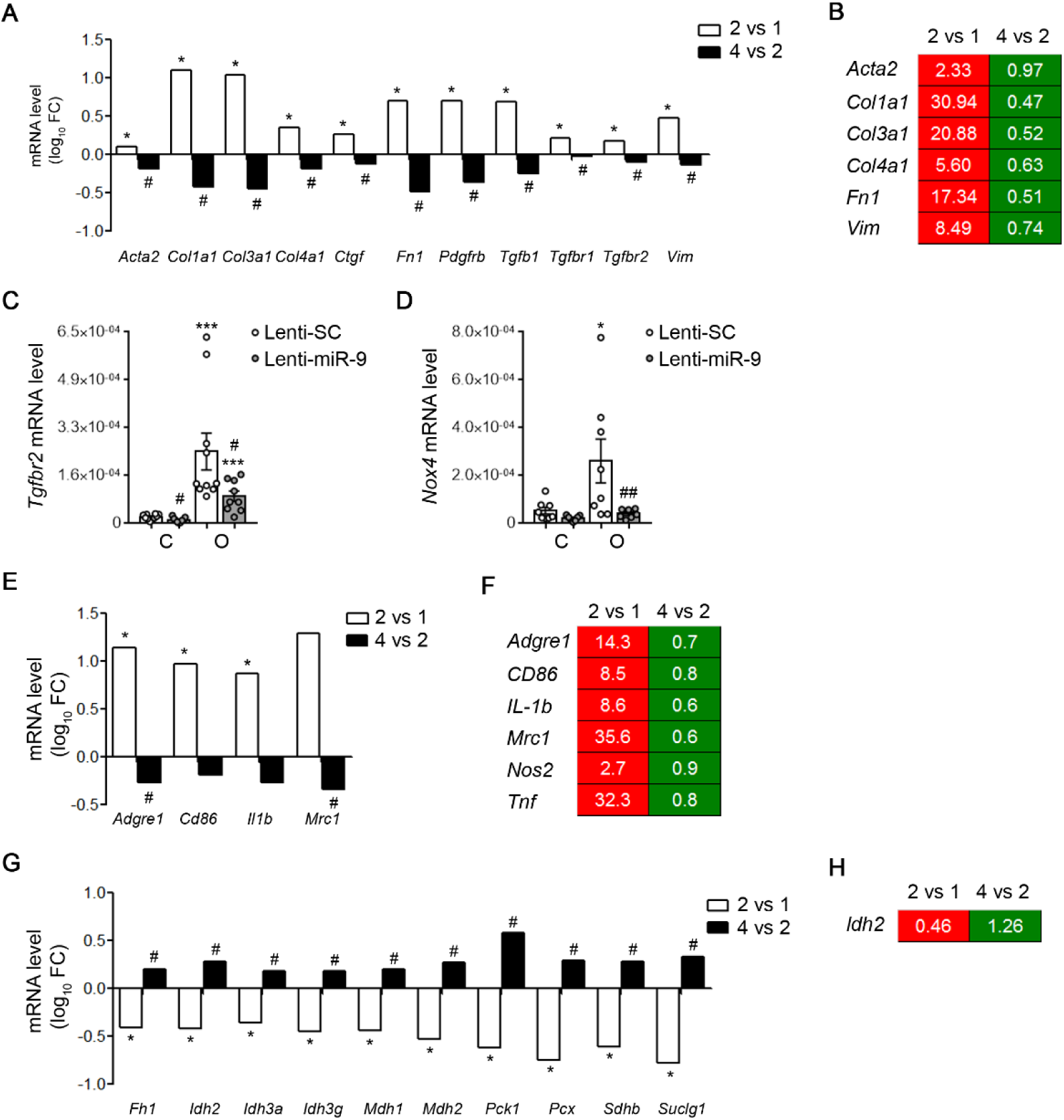
Over-expression of miR-9-5p in mice reduces the presence of fibrotic and inflammatory markers and increases the expression of tricarboxylic acid cycle (TCA)-related genes. (**A**, **B** and **E**-**H**) Analysis of gene expression in kidney samples from mice injected with lentiviral particles (2×10^7^ ifu/mouse) containing miR-9-5p (Lenti-miR-9) or scramble miRNA (Lenti-SC) 5 days before UUO. Kidney samples were analyzed 5 days after UUO. Data correspond to comparisons 2 vs 1 and 4 vs 2 (groups described in Figure 5A). (**A**, **E** and **G**) RNA-Seq data corresponding to fibrosis-(**A**), inflammatory-(**E**) and TCA cycle-(**G**) related genes. Data are represented as the log10 Fold Change (FC) (n = 3 mice per group), *Q < 0.01 compared to their corresponding contralateral kidneys; ^#^Q < 0.01 compared to obstructed kidneys in mice administered Lenti-SC, multiple hypothesis testing with the Benjamini–Hochberg FDR algorithm. (**B**, **F** and **H**) Gene expression heatmap generated by using qRT-PCR data of selected genes related to pathways described above. mRNA levels were analyzed using Taqman probes. The heatmap shows for each gene the mean of the fold change expression obtained from 6 mice per group. Analyzed genes are indicated on the left. Red indicates that the gene was down-regulated and green that the gene was up-regulated. (**C** and **D**) mRNA levels of *Tgfbr2* (**C**) and *Nox4* (**D**) miR-9-5p target genes in kidneys from mice injected as described above and subjected to UUO for 10 days were analyzed by qRT-PCR. Data are shown as the mean ± SEM (n = 7-10 mice per group), *P < 0.05, ***P < 0.001 compared to their corresponding contralateral kidneys; ^#^P < 0.05, ^##^P < 0.01 compared to obstructed kidneys in mice administered Lenti-SC, non-parametric two-tailed Mann-Whitney U test

## Supplemental Tables and Supplemental table legends

**Supplemental Table 1.** List of genes resulting from Venn intersections among mice groups described in Figure 5A. Included as a spreadsheet table.

**Supplemental Table 2.** The log10 Fold Change (FC) and their corresponding P-value of genes resulting after the scatter dot plot showed in Figure 5D. Included as a spreadsheet table.

**Supplemental Table 3.**
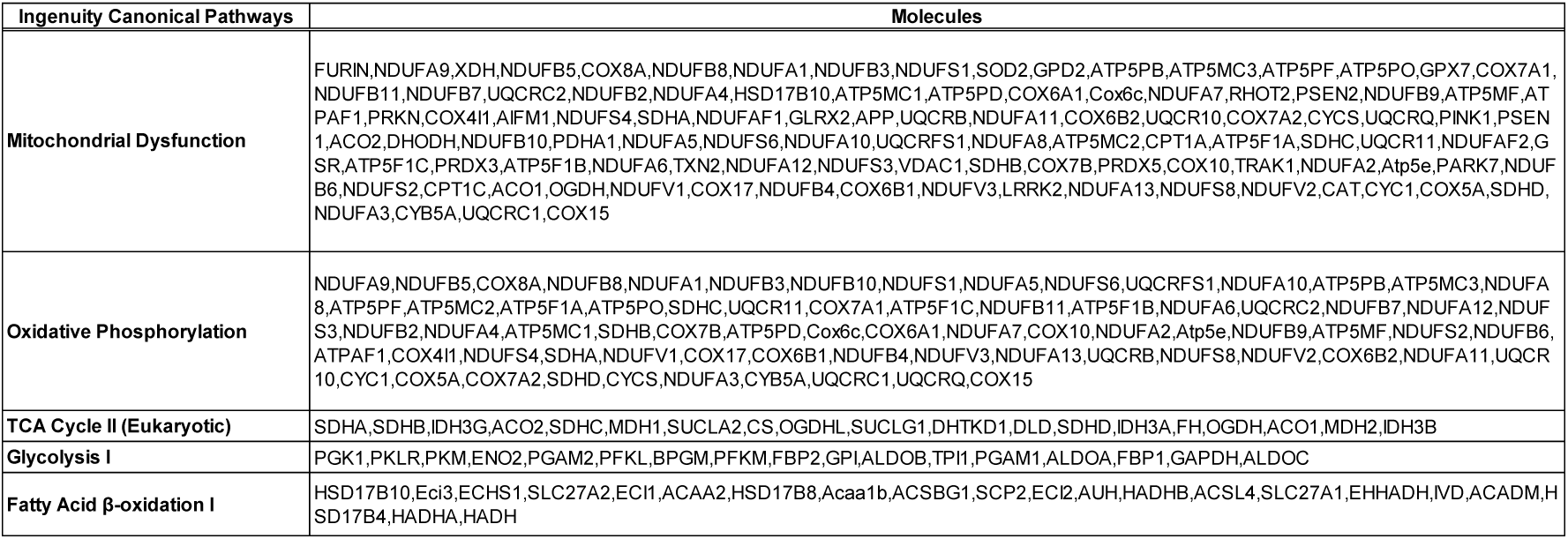
List of differentially expressed genes associated with metabolic pathways in the Ingenuity Pathway Analysis (IPA) corresponding to the comparisons between the obstructed (mice group 2) and the contralateral (mice group 1) kidneys of C57BL/6 mice injected lentiviral particles (2×10^7^ ifu/mouse) containing a scramble miRNA (Lenti-SC) and 5 days later subjected to the UUO procedure for 5 days.

**Supplemental Table 4.**
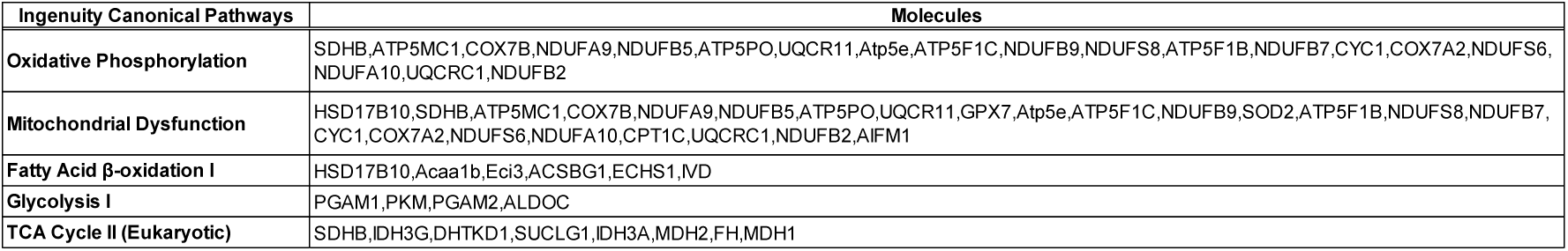
List of differentially expressed genes associated with metabolic pathways in the Ingenuity Pathway Analysis (IPA) corresponding to the comparisons between the obstructed kidneys of C57BL/6 mice injected lentiviral particles (2 × 107 ifu/mouse) containing miR-9-5p (Lenti-miR-9) (mice group 4) and obstructed kidneys of C57BL/6 mice injected lentiviral particles (2 × 107 ifu/mouse) containing a scramble miRNA (Lenti-SC) (mice group 2) and 5 days later subjected to the UUO procedure for 5 days.

**Supplemental Table 5.**
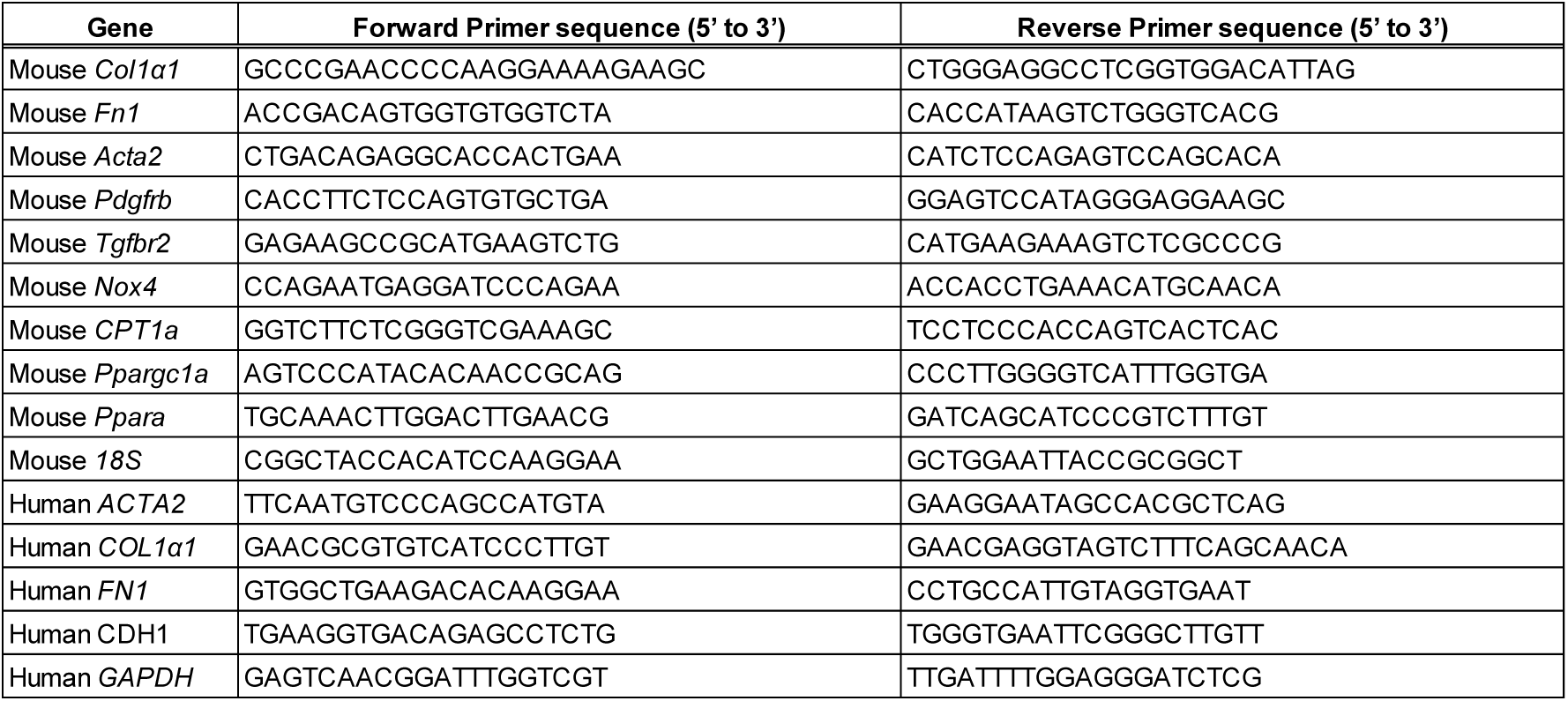
List of the primer sequences used for mRNA quantification.

**Supplemental Table 6.**
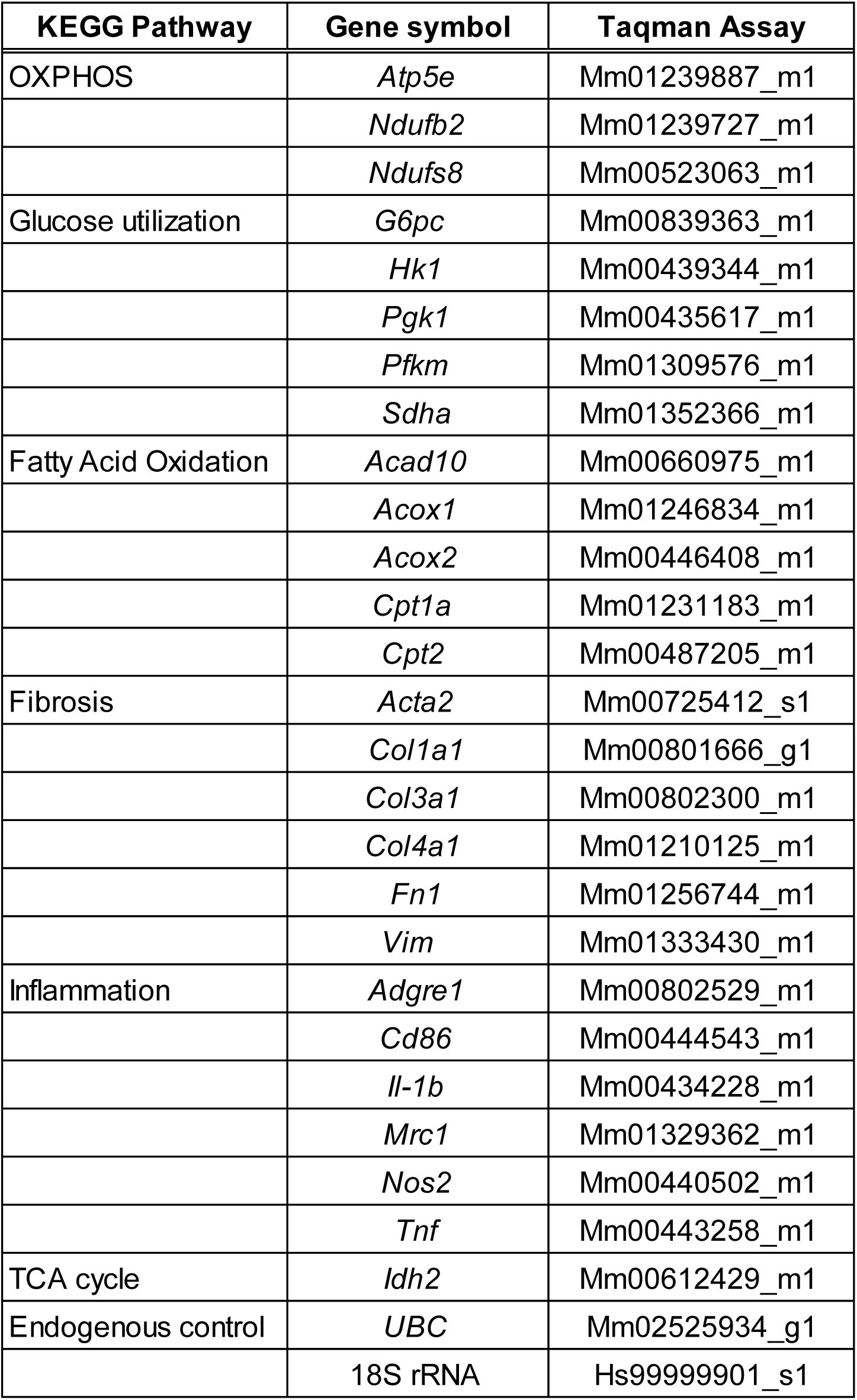
List of pathways, analyzed genes for each pathway and Taqman assay references used for mRNA quantification.

